# Phylogenomic analysis of the Lake Kronotskoe species flock of Dolly Varden charr reveals genetic and developmental signatures of sympatric radiation

**DOI:** 10.1101/2023.02.24.529919

**Authors:** Katherine C. Woronowicz, Evgeny V. Esin, Grigorii N. Markevich, Crisvely Soto Martinez, Sarah K. McMenamin, Jacob M. Daane, Matthew P. Harris, Fedor N. Shkil

## Abstract

Recent adaptive radiations provide evolutionary case studies, which provide the context to parse the relationship between genomic variation and the origins of distinct phenotypes. Sympatric radiations of the charr complex (genus *Salvelinus*) present a trove for phylogenetics as charrs have repeatedly diversified into multiple morphs with distinct feeding specializations. However, species flocks normally comprise only two to three lineages. Dolly Varden charr inhabiting Lake Kronotske represent the most extensive radiation described for the charr genus, containing at least seven lineages, each with defining morphological and ecological traits. Here, we perform the first genome-wide analysis of this species flock to parse the foundations of adaptive change. Our data support distinct, reproductively isolated lineages with little evidence of hybridization. We also find that specific selection on thyroid signaling and craniofacial genes forms a genomic basis for the radiation. Thyroid hormone is further implicated in subsequent lineage partitioning events. These results delineate a clear genetic basis for the diversification of specialized lineages, and highlight the role of developmental mechanisms in shaping the forms generated during adaptive radiation.

**Significance Statement:** Dolly Varden Charr (*Salvelinus malma*) radiation in Lake Kronotskoe provides a unique case study of the genetics of adaptation and morphological evolution. We provide first genomic and experimental analyses of this radiation and show that major axes of change may be shaped by developmental constraints.

## Introduction

The Salmonid fishes of the genus *Salvelinus* represent an exceptional example of parallel evolution and trophic adaptation to new environments (Klemetsen 2010). Charr are remarkably variable with naturally occurring, morphologically distinct populations that exhibit diverse life histories. These include complete and partial anadromy, as well as freshwater riverine and lacustrine residency (Taylor 2016, Lecaudey, Schliewen et al. 2018, Osinov, Volkov et al. 2021). Notably, freshwater resident populations frequently establish species flocks showing stereotypical morphologies, each associated with an ecological niche (Nordeng 1983, Walker, Greer et al. 1988, Sandlund, Gunnarsson et al. 1992, Maitland, Winfield et al. 2007, Simonsen, Siwertsson et al. 2017, Markevich, Esin et al. 2018, Esin, Bocharova et al. 2020, Jacobs, Carruthers et al. 2020). Even within sympatric populations, lineages with varying diets, depth preferences, and disparate spawning intervals are commonly observed (Jonsson and Jonsson 2001, Klemetsen 2010). Within independent lacustrine *Salvelinus* radiations, specific morphological adaptations have repeatedly evolved (Klemetsen 2010). Such a propensity for repeated, independent radiations make the genus *Salvelinus* an attractive system to dissect the mechanisms that facilitate rapid generation of morphological variation.

*Salvelinus* charrs have repeatedly radiated into multiple sympatric morphs with distinct evolved feeding specializations analogous to those of both African and South American cichlids (Barluenga, Stölting et al. 2006, Malinsky, Challis et al. 2015, Malinsky, Svardal et al. 2018) and African and Asian barbs (Myers 1960, Nagelkerke and Sibbing 2000, Levin, Simonov et al. 2020). Variable traits include jaw size, mouth position, eye size, pigmentation, and others. The variability within and between, charr radiations proffers important genetic case studies to uncover mechanisms of rapid morphological diversification. Such parallel events suggests that these populations experience similar selective pressures, and/or that there are genetic biases shaping the morphologies. The contributions of standing variation from ancestral populations has been suggested for other radiations, including adaptation of stickleback populations to freshwater environments (Schluter and Conte 2009). In contrast, the genetic mechanisms that enable charr to generate distinct morphologies have not yet been well defined. It has been argued that shifts in developmental timing may underlie these common transitions to specialized morphologies among charrs (Esin, Markevich et al. 2018). In this model, development biases radiations by constraining a common axis of change. Early and pleiotropic developmental shifts may give charr an exceptional capacity for adaptive radiation with only a small number of genetic modifications.

The Lake Kronotskoe radiation of Dolly Varden (*Salvelinus malma*) is unique among charrs, as it contains at least seven reproductively isolated phenotypes associating with different ecological niches (Markevich, Esin et al. 2018). This radiation is currently the broadest observed within resident lacustrine populations of this genus, and indeed, among salmonids more broadly (Markevich, Esin et al. 2018). Recent work has characterized genome-wide changes in species flocks of Alpine whitefishes (Coregonus spp.), however, those flocks comprise up to six species with varying morphologies (De-Kayne, Selz et al. 2022). Lake Kronotskoe is one of several volcanogenic lakes of the Kamchatka peninsula (**Figure 1A**) which formed approximately 12,000 years ago after a lava flow dammed the ancestral river (**Figure 1B**) (Braitseva, Melekestsev et al. 1995). The resulting landlocked Dolly Varden population diversified within the new lacustrine environment, and encompassed prominent changes in craniofacial form supporting new feeding strategies (**Figure 1C, D**) (Markevich, Esin et al. 2018, Esin, Bocharova et al. 2020). A major axis of change is proportional jaw length, seen in piscivorous Longhead and deep-water benthivorous Bigmouth morphs, as well as the modulation of frontonasal proportions found in the Nosed morphs. The deep-water omnivorous Smallmouth morph has an increase in eye size in relative proportion to the cranium as well as reduced jaw size (**Figure 1C, D**). Enlarged eye size is widespread among deep-dwelling resident charr lineages in other lakes and may suggest specialization for foraging in low light conditions (Klemetsen 2010). The varied Lake Kronotskoe charrs show strict natal homing with some morphs spawning in tributaries and other morphs spawning within the lake (Markevich, Zlenko et al. 2021). The morphs also exhibit differential annual timing windows for reproduction, which reinforces reproductive isolation and facilitates sympatric radiation (Esin, Markevich et al. 2021).

**Figure 1.**
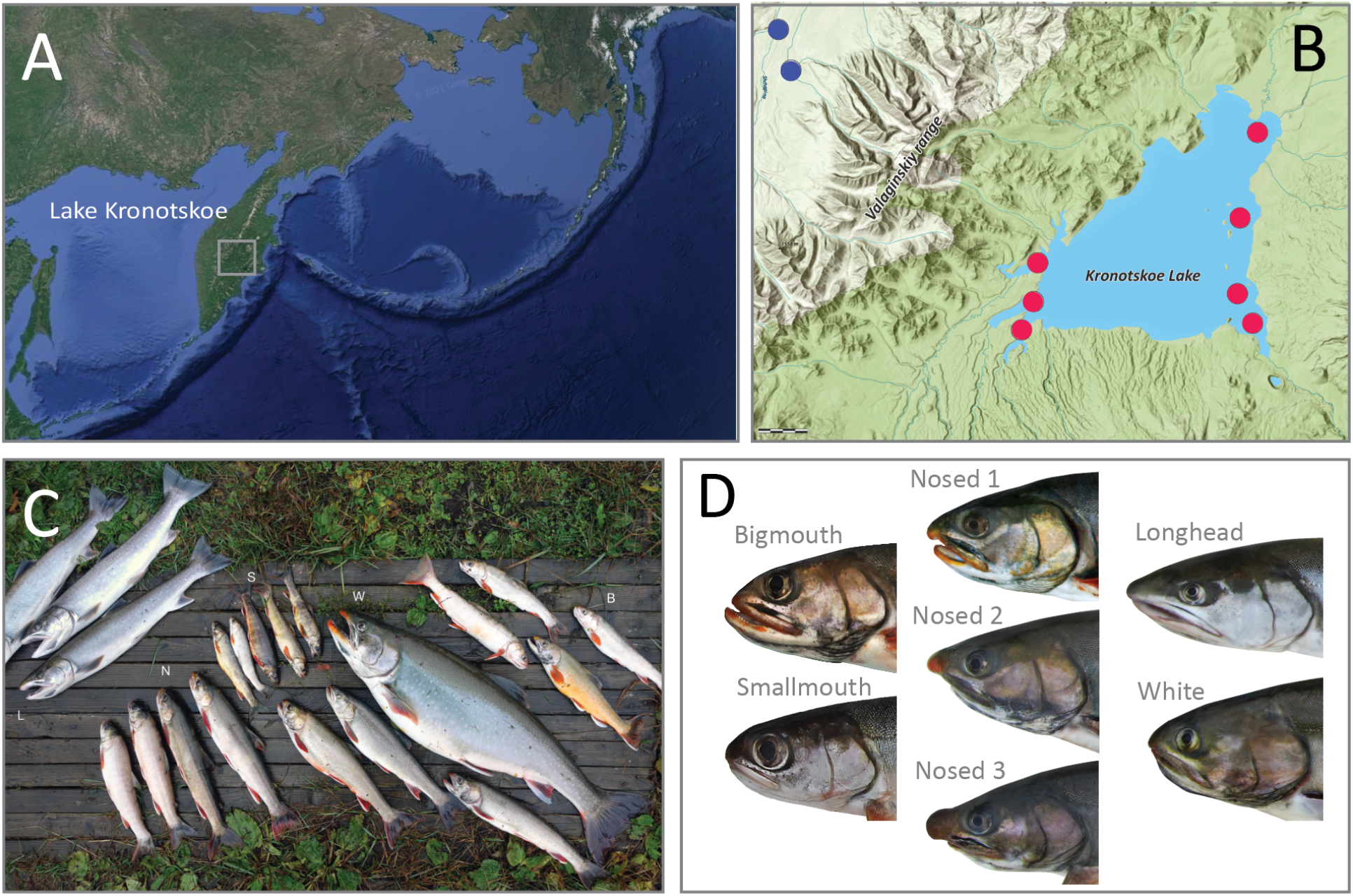
Lake Kronotskoe radiation of Dolly Varden Charr. **(A)** A map of the Kamchatka Peninsula. Lake Kronotskoe is centered within the highlighted box. **(B)** A map of Lake Kronotskoe geography and locations from which anadromous (blue dots) and resident lacustrine (red dots) Dolly Varden morphs were collected. **(C)** The Lake Kronotskoe morphs from left to right: Longhead (L), Nosed lineages (N), Smallmouth (S), White (W), Bigmouth (B). **(D)** Detailed images of representative adult heads for the seven sequenced lineages of the species flock.

Here, we employ phylogenomics to dissect the genetic context of the Dolly Varden radiation within Lake Kronotskoe. We parse the shared genetic foundation for the morphological and physiological adaptations of these specialized lineages.

### Genome wide assessment of variability within and between Lake Kronotskoe Dolly Varden morphs

To investigate variation throughout the charr genome, we performed genome-wide targeted capture of coding and non-coding loci of Dolly Varden charr from Lake Kronotskoe using a custom, pan-Salmonidae capture array. Targeted capture of small population pools allows identification of lineage-defining variation via cross-clade sequence comparisons. Similar approaches have been recently used to assess variation in Belonifomes (Daane, Blum et al. 2021), rockfishes (Treaster, Deelen et al. 2022) and notothenioids (Daane, Dornburg et al. 2019), allowing for clade-wide analysis of genomic variation and phylogenetic parsing of selective signals.

The pan-Salmonidae capture-array consists of 558,882 bait sequences designed against conserved Atlantic Salmon (*Salmo salar*) and Rainbow Trout (*Oncorhynchus mykiss*) genomic sequences and targets 97Mb of total sequence **(Figure 2 – figure supplement 1**). Coding regions constitute 83.8% of targeted regions with conserved non-coding, ultraconserved non-coding, and miRNA hairpins comprising the remaining 16.2% of targeted regions. With our capture methodology and analysis pipeline, bait sequences successfully hybridize with targeted elements harboring up to 15% deviation in sequence identity (Mason, Li et al. 2011), thereby allowing recovery and subsequent analysis of sequence variability. Pairwise analyses between homologous loci were used to distinguish fixed and variable loci between morphs. Genes associating with variation were identified by making strategic comparisons within a phylogeny focused on character diversification (Daane, Rohner et al. 2015). For outgroup analysis, we sampled anadromous Dolly Varden charr from the local Kamchatka river basin, which is adjacent to, but isolated from, that of Lake Kronotskoe (**Figure 1B**). We also sequenced a single *S. leucomaenis* individual which was collected from the same catchment as the Dolly Varden charr to serve as an outgroup.

**Figure 2.**
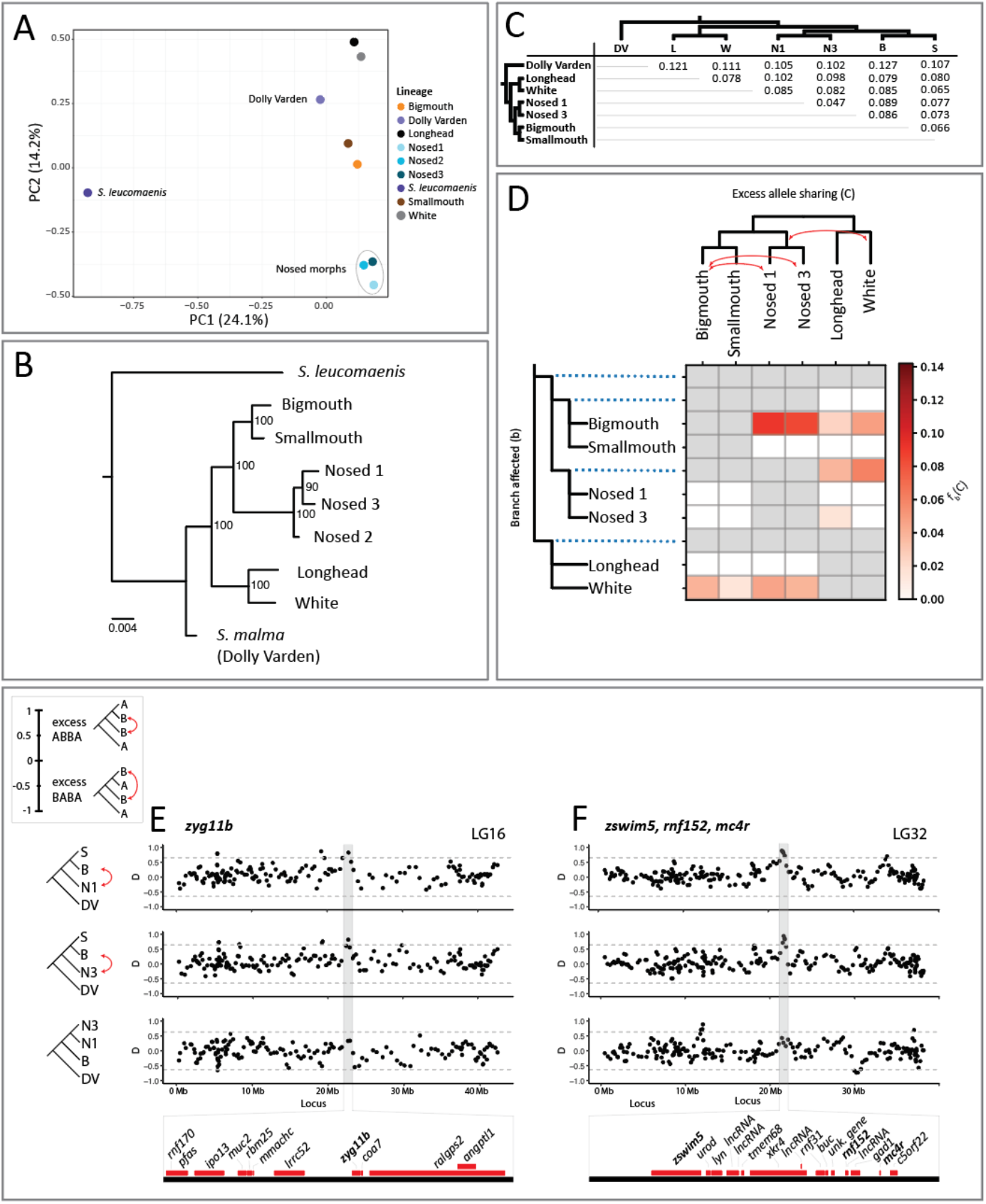
Phylogenetics and population differentiation among the sequenced lineages. **(A)** PCA plot of the seven lineages from the lacustrine species flock, and outgroups anadromous Dolly Varden and *S. leucomaenis*. PC1 distinguishes *S. Leucomaenis* from anadromous Dolly Varden and from the lake morphs. Note the tight clustering of the three Nosed morphs. **(B)** Phylogenic relationship of Lake Kronotskoe Dolly Varden lineages; anadromous *S. leucomaenis* from the Kamchatka River serve as the outgroup. **(C)** Distribution of genome-wide, pairwise Fst values calculated for non-overlapping 10kb sliding windows (**Supplementary File 5**). **(D)** The branch-specific statistic ƒ_b_ shows evidence for elevated gene flow between the Bigmouth and Nosed 1 morphs, the Bigmouth and Nosed 3 morphs, and the ancestor of the Nosed lineages and the White morph. White cells represent combinations for which p-values are not significant. Gray cells represent arrangements which are not topologically feasible for calculating ƒ-branch scores. **(E)** Sliding window plots of D for trios consisting of Smallmouth (S), Bigmouth (B), and Nosed 1 (N1), of S, B, and Nosed 3 (N3), and of N3, N1, and B. Positive D values indicate an excess of the ABBA pattern (red arrows), while negative values indicate an excess of the BABA pattern. The three plots show a common pattern of excess of allele sharing overlapping with *zyg11b* between B and N1 and B and N3, while there is no excess of allele sharing between B and N1 over B and N3. Horizontal lines signify 3SDs from the mean. Genes contained by the highlighted peak represented by red bars below plots. **(F)** three genes of interest, *zswim5*, *rnf152*, and *mc4r*, possibly constitute a supergene contained by a shared peak between B and N1 and B and N3.

For each group, we recovered approximately 90% of the targeted elements sequenced to a mean depth of 25-52 reads (**Figure 2 – Supplementary Table 1**). We detected substantial variation within each lineage as well as the putative ancestral population (avg π in Dolly Varden = 0.003) supporting pairwise analysis to detect and compare shared and unique variation within each lineage. Loci underrepresented in our capture were limited and represented as general gene classes (**Supplementary Table 2**). These elements recovered with low coverage are similar to those observed in other broad capture approaches (Daane, Dornburg et al. 2019, Daane, Blum et al. 2021, Treaster, Deelen et al. 2022).

### Genetic differentiation and relationships among Dolly Varden morphs

As a first approach, we conducted Principal Components Analysis (PCA) to visualize the relationships among our lacustrine and anadromous Dolly Varden samples and outgroup *S. leucomaenis*. PC1 separates *S. leucomaenis* from Dolly Varden and from the members of the Dolly Varden species flock (**Fig. 2A**), while PC2 broadly groups lineages according to ecological niche. Notably, all three Nosed morphs cluster quite tightly.

We investigated relationships among Lake Kronotskoe charrs through reconstruction of the phylogeny given our substantial sequence data. We used IQ-TREE to derive a phylogeny from a dataset containing 622,831 variant sites, of which 22,701 variants were informative (Nguyen, Schmidt et al. 2015). Prior interpretations of this radiation argued for multiple-step diversification based on changes in resource utilization and accompanying feeding specializations leading to the extant morphs (Markevich, Esin et al. 2018). Our phylogenomic data support the existence of the described distinct morphs, with each node having high bootstrap support (**Figure 2B**). Our analysis clusters Longhead and White morphs within a lineage that is an outgroup to the clade consisting of the Bigmouth, Smallmouth, and Nosed morphs. Nosed lineages differentiate as a distinct clade with Nosed 2 as an outgroup to Nosed lineages 1 and 3. As the Nosed 2 population associates so closely with Nosed 3 via PCA (**Figure 2A**), we focused subsequent analyses on disentangling the differentiation between Nosed morphs 1 and 3.

The anadromous and resident lacustrine Dolly Varden morphs are clearly genetically differentiated within ecologically specialized lineages, having pairwise mean Fst consistent with distinct populations (**Figure 2C**) (**Supplementary File 1**). The greatest pairwise differentiation is found between the anadromous Dolly Varden charr and lacustrine deep-water Bigmouth morph (Fst = 0.127). Notably, each lacustrine lineage is more differentiated from the anadromous Dolly Varden population than from any other lacustrine lineage. Further, lacustrine lineages are similarly differentiated from the putative ancestral riverine Dolly Varden population. Within the species-flock, the least differentiated pairing is between the lacustrine Nosed 1 and Nosed 3 morphs (Fst = 0.047) consistent with their phylogenetic relationship. Although informative of broad patterns of divergence, these values are likely underestimates of genetic differentiation due to the conserved-element-based dataset biasing analysis to regions having an inherent constraint on variation.

Introgression analysis within the clade further supports the existence of distinct lineages, although some incomplete lineage sorting was detected. Significant introgression was identified in 80% of the 20 possible trios (DSuite; Holm-Bonferoni FWER < 0.01) in patterns which deviated from the topology of the phylogeny (**Supplementary Table 3**) (Malinsky, Matschiner et al. 2021). Despite observed significance, the introgression values were relatively small compared to the recent timeline of the Lake Kronotskoe radiation. The Bigmouth and Nosed 1 morphs showed the greatest excess of allele sharing (D_tree_ = 4.2%) (**Supplementary Table 3**). We further calculated ƒ_4_-admixture ratios and used ƒ-branch statistics to disentangle the interrelatedness of admixture signals among morphs that share common internal phylogenetic branches (**Figure 2D**). The greatest proportions of shared alleles were between the Bigmouth morph and the Nosed 1 morph (ƒ-branch = 9.0%), between the Bigmouth morph and the Nosed 3 morph (ƒ-branch = 8.3%), and between the White morph and the ancestor of the Nosed 1 and Nosed 3 morphs (ƒ-branch = 5.9%). The ƒ-branch statistic further supported the interpretation that Lake Kronotskoe lineages differentiated while maintaining relative genetic isolation.

To determine the extent to which Nosed 1 and Nosed 3 share common introgressed loci with Bigmouth, we calculated D in sliding windows (40 variant windows, 20 variants step-size) for Smallmouth, Bigmouth, Nosed 1 (mean = 0.046, SD = 0.22) and Smallmouth, Bigmouth, Nosed 3 (mean = 0.041, SD = 0.21). Anadromous Dolly Varden served as the outgroup for all analyses of introgression. Among sliding windows with D >= 0.8 (32 of 10,617 windows for S, B, N1; 28 of 10,790 windows for S, B, N3), we observed shared, isolated regions of the genome with signatures of admixture (**Supplementary File 2**). Among these, we detected a large interval with evidence for shared introgression between Bigmouth and Nosed morphs 1 and 3 (mean D for the interval = 0.53 for S, B, N1; 0.54 for S, B, N3) with of more than one third of windows with D > 3 SDs from the mean (**Figure 2 – figure supplement 2**). However, most introgressed intervals were more spatially restricted. One locus potentially associating with craniofacial differentiation spanned an interval centered on *zyg11b* which has been implicated in craniofacial microsomia (**Figure 2E**)(Tingaud-Sequeira, Trimouille et al. 2020). Another locus included multiple genes of interest including, including *zswim*5, which is expressed in cranial neural crest (Wong, Rebbert et al. 2016), *rnf152*, which is involved in regulating neural crest formation (Yoon, Kim et al. 2022), and *mc4r*, which is a crucial regulator of appetite and metabolism via leptin and thyroid signaling modulation (**Figure 2F**)(Decherf, Seugnet et al. 2010). This suite of craniofacial and metabolic genes may have functioned like a superlocus which was spread via shared introgression between the Bigmouth and Nosed lineages and subsequently reinforced the differentiation of distinct lineages.

### Patterns of differentiation between river and lake populations of Dolly Varden charr

To understand shared variation that differentiates the Lake Kronotskoe residents from the river-caught anadromous population of Dolly Varden, pairwise Fst was calculated per coding (25,373 genes) and conserved non-coding elements (CNE)(22,575 CNEs) from our targeted capture (**Figure 3A, B**). Fst values were computed by grouping all variation contained within the species flock and comparing against the variation contained in the riverine Dolly Varden charr population (lake versus river). There are 327 genes (**Supplementary File 3**) and 80 CNEs (**Supplementary File 4**) with Fst > 0.5. Each CNE was assigned to flanking target genes by GREAT (McLean, Bristor et al. 2010). Some intervals harbor multiple highly differentiating CNEs. For example, the close proximity of two CNEs led to two hits for *cholinesterase-like* and *dync2h1* (**Figure 3B**). Interestingly, craniofacial (orange) and thyroid (blue) related genes (5 craniofacial genes of top 20, 2 thyroid genes of top 20) and CNEs (12 craniofacial CNEs of top 20, 3 thyroid CNEs of top 20) are enriched across the most differentiating elements (**Figure 3A, B**).

**Figure 3.**
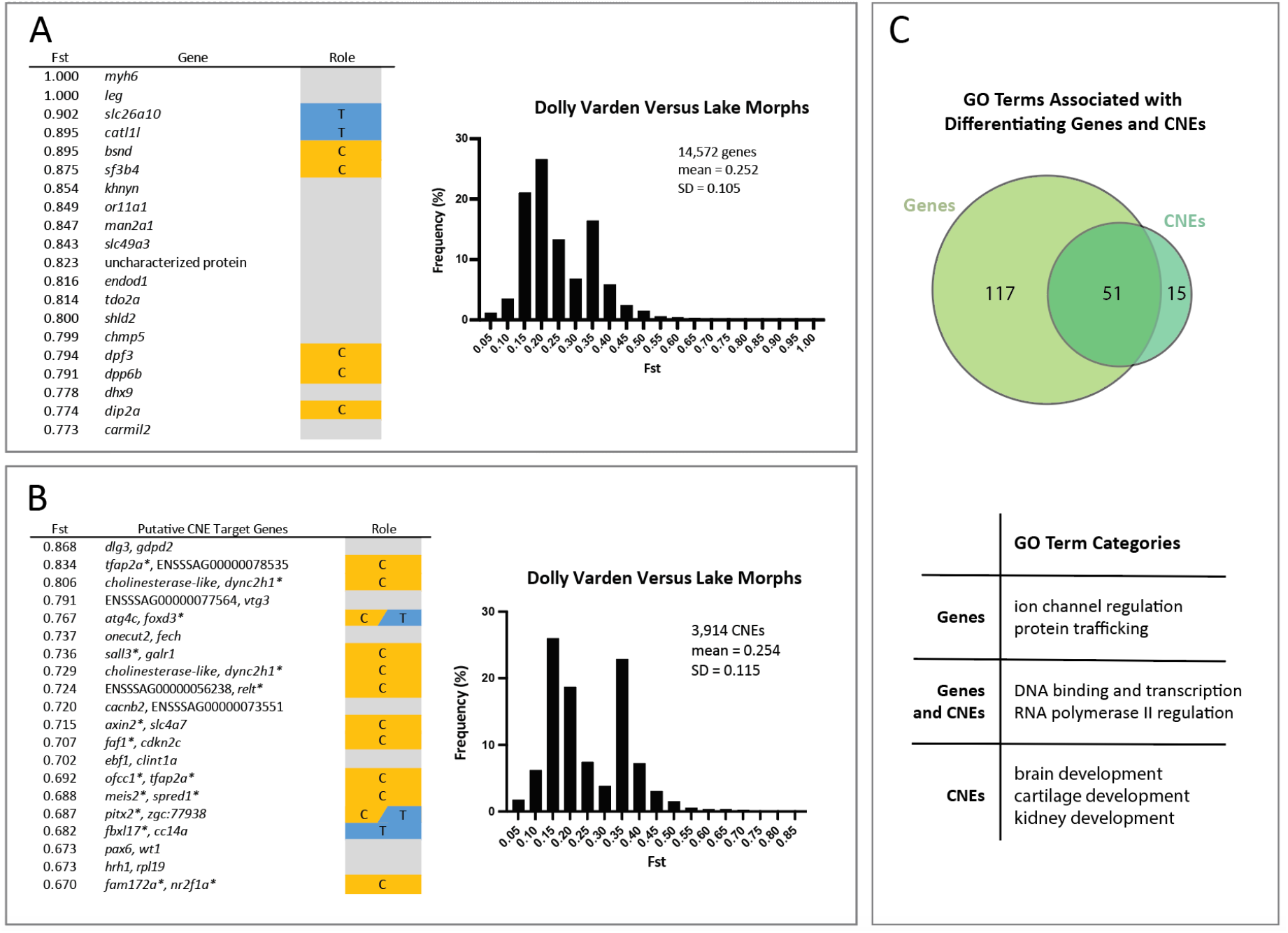
Coding and non-coding conserved elements differentiating anadromous Dolly Varden charr from the Lake Kronotskoe species flock. **A**) Table showing the top 20 genes differentiating riverine Dolly Varden from Lake Kronotskoe inhabitants along with distribution of Ft values of genes. B) Table of the top 20 differentiating CNEs and the distribution of Fst values. C) Overview of shared and unique GO terms associated with genes and CNEs with Fst > 0.5. Venn diagram values represent GO terms which occur six or more times. Table summarizes broad categories of most frequent GO term associations. The distributions and the numbers of elements represent all non-zero Fst values; *orange*, genes with known roles in modulating craniofacial morphology (C); *blue*, genes with known roles in thyroid function (T). Asterisk denotes gene classified as (C), (T), or both.

Craniofacial genes assigned to highly differentiating CNEs by GREAT analyses included both those with predicted roles in neural crest (*tfap2a*) (Rothstein and Simoes-Costa 2020), as well as frontonasal (*meis2, pitx2*) (Evans and Gage 2005, Fabik, Kovacova et al. 2020) and splanchnocranium (*faf1*) development (Ma, Zhu et al. 2017). Of note, *pitx2* is also involved in release of thyroid stimulating hormone from the pituitary (Castinetti, Brinkmeier et al. 2011). *dync2h1* and *tfap2a* appeared more than once in our analyses as putative regulated loci, suggesting compounding alterations to the regulatory environment at these loci may contribute to the genetic landscape that distinguishes the riverine Dolly Varden from the lacustrine morphs. Some CNEs are flanked on both sides by genes implicated in craniofacial development (*ofcc1* and *tfap2a*, *meis2* and *spred1*, *fam172a* and *nr2f1a*). Genes connected to ion transport (*myh6, khnyn, or11a1, endod1, cacnb2,* and *hrh1*), and vesicular transport (*man2a1, shld2, gdpd2, onecut2,* and *clint1a*) according to KEGG and Reactome reconstructions were also well represented among the top differentiating genes and as targets of the top differentiating CNEs. The majority of nucleotide changes identified were single SNPs with unknown functional significance. However, high Fst is reflective of potential selection or bottleneck at the locus.

GO terms were assigned to all genes (2,825 GO terms) and CNEs (1,419 GO terms) that differentiate anadromous Dolly Varden from Lake Kronotskoe residents (Fst > 0.5). Among GO terms appearing 6 or more times, specific themes emerged (**Fig. 3C**). Highly differentiating genes are associated with roles in ion channel regulation and protein trafficking. Highly differentiating CNEs are associated with brain, kidney, and cartilage development. Other GO terms shared among highly differentiating coding and noncoding elements include functions such as DNA binding, transcription factors, and regulation of RNA polymerase II (**Supplementary file 5**).

Within the candidate genes, there are signals related to regulation of thyroid signaling in development. For example, the gene *slc26a10,* encodes a sulfate transporter that functions in thyroid hormone synthesis and also acts downstream of Thyroid hormone receptor alpha (THRa) (Richard, Guyot et al. 2020). We found that *slc26a10* has segregating nonsynonymous changes in conserved residues that are highly differentiated between riverine and lacustrine morphs (**Figure 4A, B**). However, the function of this gene is not well understood, and alignment-based analyses predict this amino acid substitution to have little effect on function (neutral by PROVEAN, tolerated by SIFT) (Ng and Henikoff 2003, Choi and Chan 2015). By Fst score, the next candidate gene, *catl1l,* encodes Cathepsin L1-like and is orthologous to Cathepsin L1. This gene may be involved in the processing of thyroid prohormone (Friedrichs, Tepel et al. 2003). Next, splicing factor 3b subunit 4 (*sf3b4)* is a key developmental regulator of frontonasal patterning and growth and is associated with craniofacial Nager syndrome in humans (Bernier, Caluseriu et al. 2012, Petit, Escande et al. 2014). Within this locus, we find synonymous variants at high Fst in all lacustrine morphs, and fixed within benthic groups (**Figure 5A, B**). These elevated footprints of clustered, highly differentiating variants may indicate the presence of further modifications within poorly conserved non-coding regions in linkage to these sites which were not targeted by our bait sequences.

**Figure 4.**
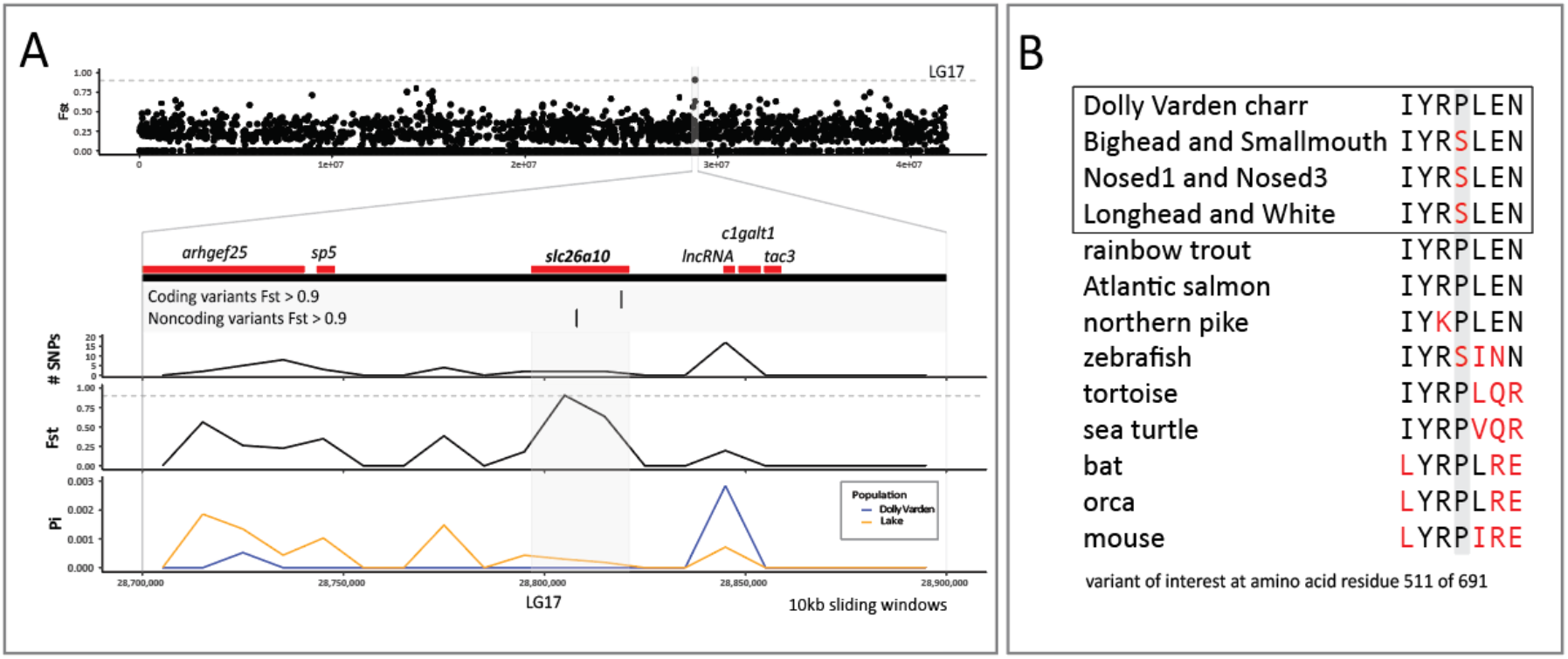
Differentiation of *slc26a10* in pairwise comparisons between anadromous Dolly Varden charr and the Lake Kronotskoe species flock. **(A)** The *slc26a10* locus shows high differentiation (Fst). The gene locus contains one highly differentiating non-coding and one coding variant. Dolly Varden *slc26a10* is homologous to human pendrin (*SLC26A4*), a known thyroid regulator. The broader locus has low nucleotide diversity as illustrated by sliding window plots of Tajima’s Pi. Horizontal dotted gray line represents Fst = 0.9. Plots are included for 10kb sliding windows along LG17, the coding elements within the broader locus, the number of variants per sliding window, Fst, and Tajima’s Pi per sliding window in pairwise comparisons between riverine Dolly Varden charr and the Lake Kronotskoe species flock. **(B)** Slc26a10 contains a fixed amino acid substitution in a conserved proline that differentiates Dolly Varden charr from each of the major clades of the lacustrine morphs.

**Figure 5.**
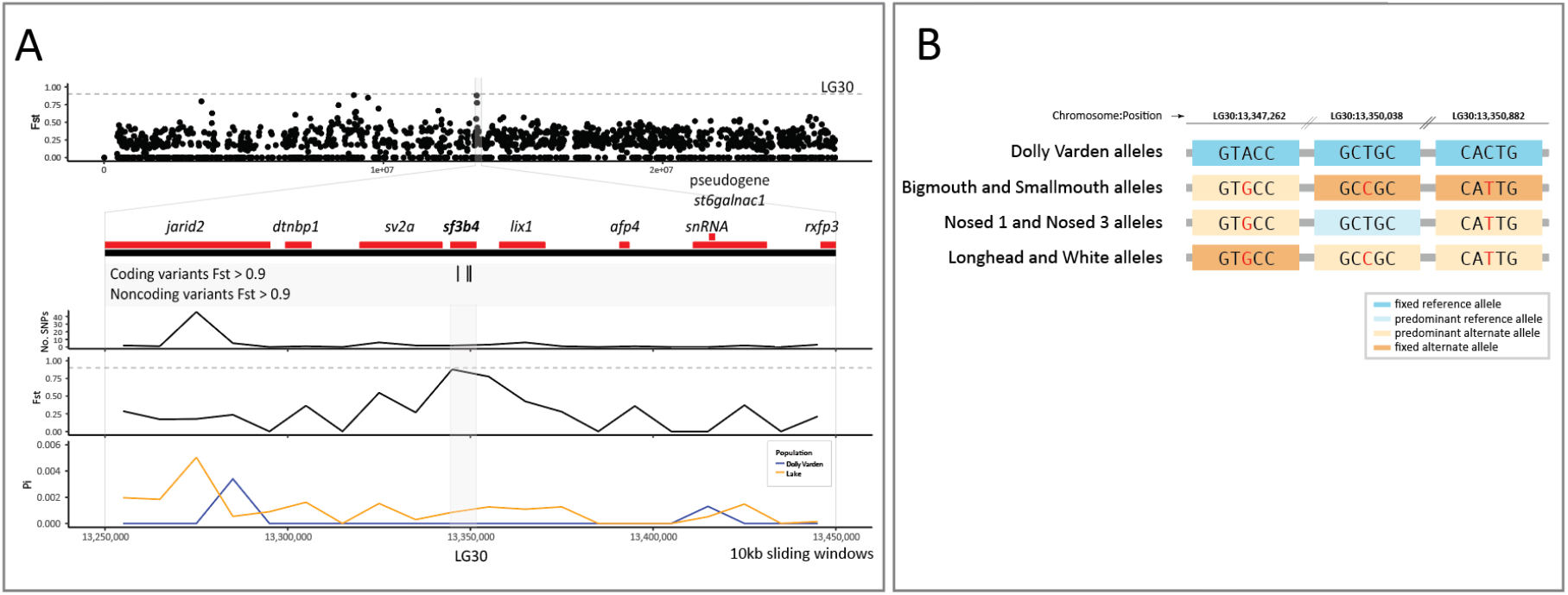
Fixation of variation in s*f3b4* between anadromous Dolly Varden charr and the Lake Kronotskoe species flock. **(A)** Three highly differentiating synonymous variants lie within *sf3b4*. Fst is plotted in non-overlapping 10kb sliding windows along the length of the chromosome LG30. *Sf3b4* is contained within a 200kb interval of low nucleotide diversity as illustrated by sliding window plots of Tajima’s Pi. Horizontal dotted gray line represents Fst = 0.9. The broad locus encompassing *sf3b4* is shown in detail including plots of the number of variants, Fst, and Tajima’s Pi in 10kb non-overlapping sliding windows. **(B)** 3 synonymous variants in *sf3b4* are fixed in lacustrine Dolly Varden charr; *dark orange,* non-reference allele fixed*; light orange,* non-reference allele predominant (alt. allele freq. > 50%) in sequence pool; *Light blue*, a lineage for which the reference allele is predominant (ref. allele freq. > 50%) at the locus.

### Thyroid hormone signaling activity is associated with shifts in craniofacial form

As modulation of thyroid signaling has been implicated in phenotypic specialization of charrs in different environments (Esin, Markevich et al. 2021), the pattern of fixation in thyroid-associated genes and non-coding elements in lacustrine morphs compared with riverine Dolly Varden warranted further investigation. To address if changes in thyroid metabolism are associated with different morphs in Lake Kronotskoe, we assessed levels of circulating thyroid hormone in adult riverine and lacustrine charr individuals from the Lake. Intriguingly, we found a significant decrease in T_3_ (the most genomically active form of thyroid hormone) hormone in specific lacustrine populations (**Figure 6A**). The pattern of reduced T_3_ abundance across the species flock correlates with a clear change in craniofacial proportions: Nosed and Smallmouth morphs, with sub-terminally positioned mouths, have significantly decreased T_3_ levels. This is in stark contrast to riverine Dolly Varden charr and the lacustrine White morph, which have comparatively ‘wild-type’ craniofacial form, and the piscivorous Longhead and deep-water benthivorous Bigmouth morphs, which are highly specialized (**Figure 1D**).

**Figure 6.**
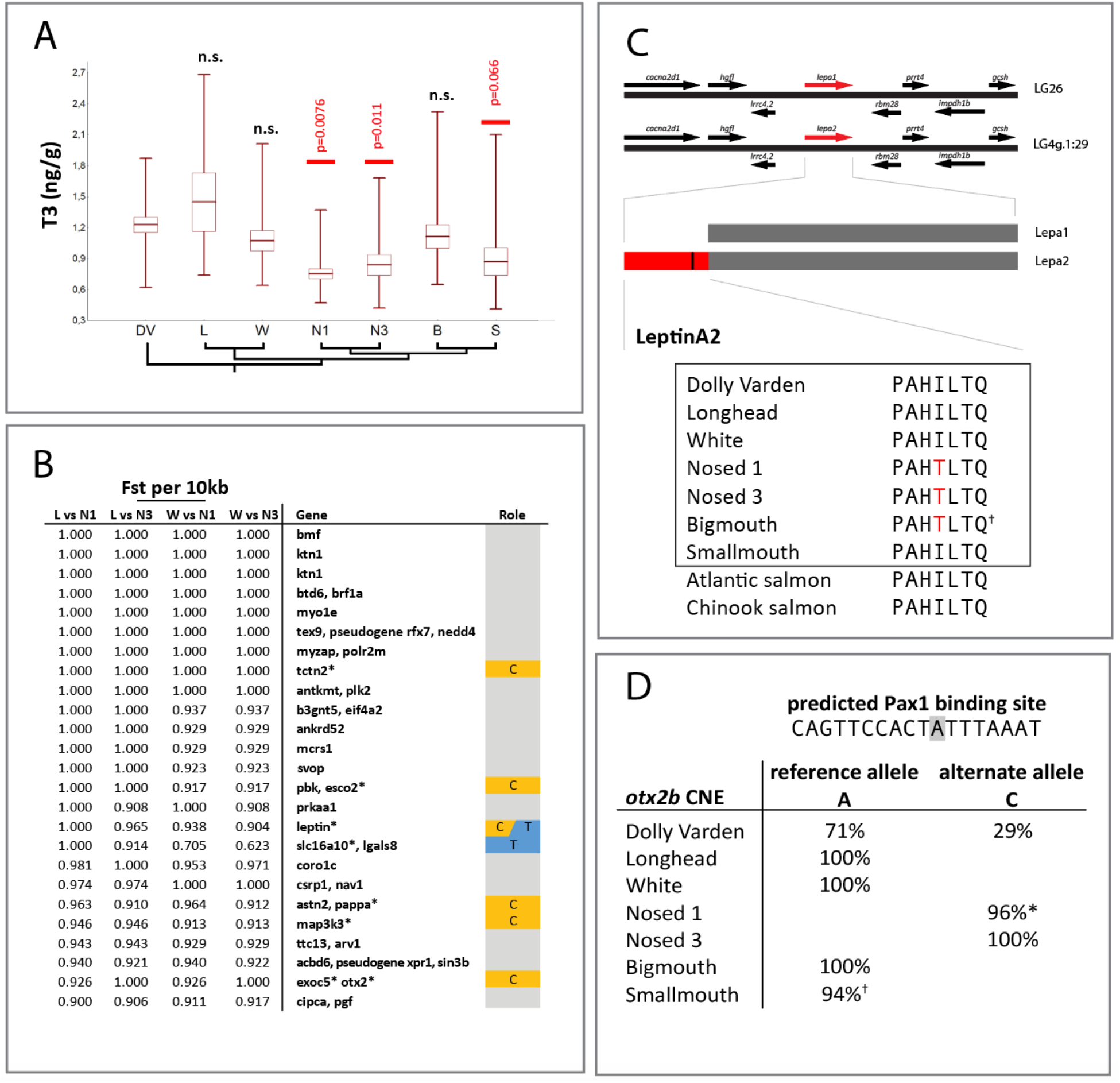
Thyroid hormone (T3) levels and associating highly differentiating candidates in lacustrine Dolly Varden morphs related to thyroid function and craniofacial morphology. **(A)** Serum T3 levels were significantly less in Nosed 1, Nosed 3, and Smallmouth lineages relative to anadromous Dolly Varden; mean +/- 1 SD, with boundary values indicated (min/max). **(B)** Top 20 candidate regions differentiating Nosed lineages from Longhead and White lineages. Sites uniquely differentiating Longhead morphs from Nosed 1 and Nosed 3 morphs, and White morphs from Nosed 1 and Nosed 3 morphs identify loci associated with thyroid function (blue, T) or craniofacial developmental (orange, C) modulating elements. **(C)** Schematic of Leptin A paralogs in salmonids detailing previously unknown gene *leptin a2* having a unique 66 amino acid N-terminus extension compared to its paralog. Nosed lineages contain a fixed, non-synonymous SNP in this conserved N-terminal sequence. (**D**) An *otx2b* CNE contains fixed or highly differentiating variants within a predicted Pax1 binding domain that associates with lake morphs exhibiting significantly different thyroid signaling activities; percentage of reference and alternate allele reads indicated per morph. Low level detection of variant or reference alleles noted (1 read for reference allele (asterisk), 2 reads for non-reference allele (dagger)).

Our findings identify a disproportionate number of differentiated loci in Nosed lineages that are known to regulate thyroid signaling (**Fig. 6B**). We assessed specific mutations identified as fixed or nearly fixed (Fst > 0.9) within Nosed morphs that differentiate them from Longhead or White morphs. A candidate locus that differentiates the low T_3_ Nosed 1 and Nosed 3 morphs from the Longhead and White outgroups is a *leptin* homolog (**Fig. 6C**). All salmonids have two leptin A ohnologs derived from the shared salmonid whole genome duplication. LeptinA2 paralogs encode an N-terminal sequence extended by 66 amino acids compared to LeptinA1 or B orthologs within salmonids. The nonsynonymous change differentiating Nosed lineages lies within a conserved residue of the unique sequence of this leptin paralog. As leptin reciprocally modulates thyroid hormone activity, both endocrine signaling pathways affect global metabolic activity. Notably, *mc4r*, which shows evidence of allele sharing between the Bigmouth and Nosed lineages, serves as a relay through which leptin stimulates the thyroid axis, suggesting that these hormonal axes may be modulated at multiple levels in differentiated lineages (Decherf, Seugnet et al. 2010). Exogenous leptin and thyroid hormone treatments are associated with enlarged craniofacial dimensions marking an intriguing relationship between these pathways and Nosed lineage diversification (Yagasaki, Yamaguchi et al. 2003, Zimmermann-Belsing, Brabant et al. 2003, Copeland, Duff et al. 2011, Shkil, Kapitanova et al. 2012, Keer, Cohen et al. 2019).

We also identified a highly differentiated variant within a conserved non-coding enhancer of *otx2b* (**Fig. 6D**). Nosed 1 and Nosed 3 morphs are fixed or nearly fixed for a variant allele which lies in a predicted *Pax1* transcription factor binding site. *otx2b* is involved in development of the skull and the anterior pituitary gland, which regulates hormonal signaling including the thyroid axis (Diaczok, Romero et al. 2008, Bando, Gergics et al. 2020). Due to the significant overlapping domains of *pax1* and *otx2* expression in the pharyngeal arches and the oral endoderm from which the pituitary gland arises, the differentiating variant identified provides a plausible regulatory shift associated with evolution of craniofacial morphology (Liu, Wang et al. 2013, Liu, Lin et al. 2020).

We asked whether the subterminal mouth positions exhibited by the low-T_3_ lineages (Smallmouth, Nosed 1, and Nosed 3) could be caused by the reduced plasma thyroid hormone levels in these morphs. We used transgenic thyroid ablation to determine whether experimental hypothyroidism (McMenamin, Bain et al. 2014) caused any parallel shifts in subterminal mouth position in the zebrafish system. Indeed, hypothyroid zebrafish showed a significant shift in the maxilla position, moving ventrally from a supra-terminal position (**Fig. 7A, B**). Further, to test whether the top candidate genes were altered in a hypothyroid context, we extracted mRNA from the heads of control and hypothyroid larval zebrafish at 7 and 14dpf and quantified expression levels by RT-qPCR (**Fig. 7C**). While there is strong genetic differentiation between the lake morphs and the riverine Dolly Varden for *slc26a10* and *sf3b4*, we did not detect any significant difference in gene expression levels. We also quantified gene expression for *otx2b* and its putative regulators, *pax1a* and *pax1b*. While genes differentiated the Nosed morphs, we did not detect significant differences in gene expression levels in the head. However, *lepa* was significantly upregulated in the heads of 14dpf hypothyroid larvae, indicating that under normal developmental conditions, thyroid hormone suppresses *lepa* expression in the head.

**Figure 7.**
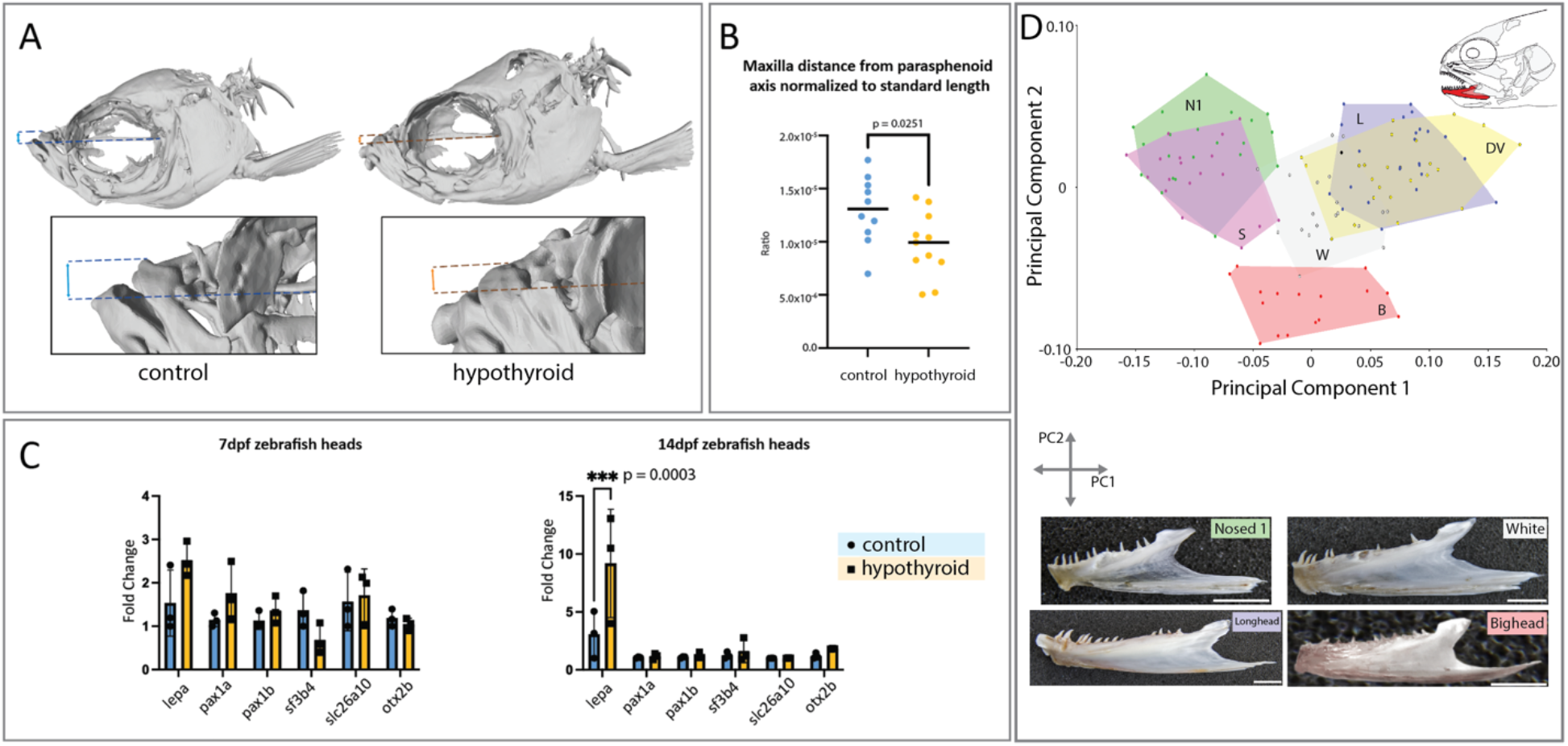
Effect of thyroid follicle ablation upon morphology and gene expression. **(A)** In thyroid ablated adult zebrafish (hypothyroid) there is a significant shift in the position of the maxilla towards a more terminal position. **(B)** In hypothyroid individuals, there is a significant reduction in the distance from the dorsal-most position of the maxilla to the long-axis of the body. All distances are relative to the standard length. n = 11 for each condition. **(C)** In hypothyroid individuals, there is a significant increase of lepa expression in the head at 14dpf. N = 3 pools of 20 heads for each condition. **(D)** Plotting Principal Components 1 and 2 reveals significant differences in dentary shape (Proscrustes ANOVA F_40;944_=40.84 p˂0.0001) between morphs. Scale bar = 10mm.

Work in zebrafish previously identified skeletal elements that are phenotypically sensitive to thyroid hormone titer (Keer, Cohen et al. 2019, Keer, Storch et al. 2022); we used geometric morphometrics to evaluate these skeletal elements in charrs. Among charr lineages, we compared the shapes of the dentary, anguloarticulare, hyomandibula and parasphenoid. Mirroring the pattern of zebrafish, charrs significantly differ in the shape of TH-sensitive bones (Procrustes Anova for dentary and anguloarticulare F_40;944_=40.84 p˂0.0001; F_80;1760_=18.88 p˂0.0001, respectively) and display subtle differences in the shape of TH-insensitive bone (Procrustes Anova for hyomandibula F_60;1212_=6.55 p˂0.001)(**Figure 7D, Figure 7 – figure supplements 2, 3**). The parasphenoid, which forms the neurocranial base, is not variable (**Figure 7 – figure supplement 4**). These data were supported by pairwise calculation of Procrustes distances between morphs, which demonstrate significant differences (p˂0.001) in the shape of jaw bones between all morphs, excluding pairs: L-DV and N1-S, and absence of differences in the shape of parasphenoid between most of morphs, excluding pairs formed by B, S and piscivorous morphs (L and W). Performing principal component analysis (PCA) on the dentary and anguloarticulare revealed that the distribution of morphs along PC1 (a component explaining 66.9% and 57.7% of the variance, respectively) (**Fig. 7B**) corresponded to their distribution along the T_3_ value axis (**Fig. 6A**).

## Discussion

Lake Kronotskoe harbors a unique radiation of Dolly Varden charr that provides a powerful new case to study the genetic and developmental foundations supporting vertebrate radiations. Previous models centered on morphology, ecology and feeding behavior suggest two lacustrine clades: a deep-water clade consisting of Smallmouth and Bigmouth morphs, and a shallow-water (pelagic and littoral) clade composed of Longhead, White, and Nosed morphs (Markevich, Esin et al. 2018). Our data redefine the evolutionary relationships among the lake morphs and support differentiated true-breeding lineages.

Lake Kronotskoe arose from a volcanogenic event and presently drains via a waterfall considered impassable by charr (Viktorovskii 1978). Pairwise Fst analyses found that each lake lineage is more differentiated from riverine Dolly Varden than from any other lineage. Furthermore, each lake lineage is differentiated from riverine Dolly Varden charr at roughly equivalent levels. We determined that the Lake Kronotskoe lineages maintain reproductive isolation through low hybridization. By contrast, the much older Lake Malawi cichlid radiation, has ƒ-branch values commonly exceeding 5% and up to 14.2% (Malinsky, Svardal et al. 2018) and *Coregonus* salmonids also have extensive introgression (De-Kayne, Selz et al. 2022). These new data suggest that the species flock in Lake Kronotskoe was established by a single founding population having shared genetic signatures that quickly established and maintained thoroughly reproductively isolated populations.

We found specific genetic differentiation between lacustrine resident lineages and riverine Dolly Varden populations with selective signatures in genes regulating craniofacial development and thyroid function. It is important to note that while PROVEAN and SIFT predicted the serine substitution shared among the lake morphs would have little consequence upon *slc26a10* function, such alignment-based prediction methods tend to underestimate potential effects if variation is shared in other lineages (Ng and Henikoff 2003, Choi and Chan 2015). Indeed, the multiple sequence alignment showed that zebrafish also encode serine at that residue.

While the relationship to morphological radiation is less clear, the abundance of ion transport and protein trafficking genes among the highly differentiating genes, suggests that these processes may also be important drivers of charr evolution. Crucially, ion homeostasis is central to effective osmoregulation during freshwater adaptation (McCormick, Regish et al. 2019). Proper ion channel expression is also a factor in chondrocyte maturation and homeostasis (Dicks, Maksaev et al. 2023, Brylka, Alimy et al. 2024). Whether and how protein trafficking contributes to adaptations in Lake Kronotskoe is less obvious.

The data show that CNEs are particularly influential in the evolution of kidney function and cartilage morphology within the Lake Kronotskoe radiation. Differentiating CNEs are associated with development of the brain, kidneys, and cartilage. The kidneys are indispensable for osmoregulation and obligate freshwater populations of salmonids place different demands on their kidneys their anadromous counterparts (Tipsmark, Sørensen et al. 2010). Cartilage templates lay foundations for many craniofacial structures. In concert, variation of these three traits may contribute to differential behavior, physiology, and morphology characterized within this adaptive radiation.

Genes that function in thyroid hormone regulation are differentially selected, suggesting modulation of the hormone may underlie stereotypical phenotypic shifts in lacustrine charr morphology. As many specialized morphologies are hypothesized to arise from heterochronic shifts in development (Simonsen, Siwertsson et al. 2017), the thyroid axis may prove to be a common mechanism underlying the adaptive potential and may contribute to the remarkable similarity of morphologies exhibited by lacustrine charr species flocks (Esin, Markevich et al. 2021, Esin, Markevich et al. 2021). Indeed, the pairs of morphs in each of the Lake Kronotskoe clades exhibit alternative heterochronic tendencies, which could result from alterations in thyroid signaling. For example, the Smallmouth morph with proportionally large eyes and blunt, rounded rostra shows hallmarks of paedomorphosis, while the sister Bigmouth morph possesses peramorphic traits such as overdeveloped lower jaw. The enlarged jaws and frontonasal protrusion of the Longhead morph, as well as the drastically modulated frontonasal proportion of the Nosed 3 morph are peramorphic features in comparison to their sister lineages. Similar TH-induced craniofacial changes also arose during the radiation of charrs dwelling in other lakes and rivers. For example, in Lake El’gygytgyn, there resides an extremely low TH-content small-mouth charr (*S. elgyticus*) has big eyes and blunt, rounded rostra, while the closely related boganida charr (*S. boganidae*) with a high TH-content has elongated jaws (Esin, Markevich et al. 2021, Esin, Shkil et al. 2024). The piscivorous stone charr *S. malma* lineage, dwelling in sympatry with typical Dolly Varden the Kamchatka river and characterized by a high TH-level, displays an accelerated rate of ossification of the tooth-bearing bones, reduced eye size, elongated head, and big mouth as its definitive morphological traits (Esin, Markevich et al. 2020).

Indeed, thyroid hormone-induced adaptive morphologies are found in phylogenetically distant fishes, the large African barbs (g. *Labeobarbus*; Cypriniformes; Teleostei), inhabiting Lake Tana (Nagelkerke and Sibbing 2000). The age of the Lake Tana species flock of barbs is comparable with the Lake Kronotskoe species flock, yet genetic differences between Lake Tana morphs are comparatively subtle (de Graaf, Megens et al. 2010, Nagelkerke, Leon-Kloosterziel et al. 2015), and ecomorphological differentiation is a result of heterochronic shifts presumably induced by thyroid axis alterations (Shkil, Lazebnyi et al. 2015). Such similarities suggest that genetic modification of thyroid signaling may be a widespread mechanism facilitating rapid freshwater teleost adaptive radiations; thyroid modifications could provide a pleiotropic foundation from which more specialized morphologies may be further elaborated. Our experimentally induced hypothyroidism lend support to this possibility: ablating thyroid follicles in the zebrafish creates a shift towards a subterminal mouth position, recapitulating the morphology of the low-T3 charr lineages. Further, the craniofacial elements variable among the Lake Kronotske charrs are the same bones that are known to be sensitive to thyroid hormone alterations in a zebrafish context (Keer, Cohen et al. 2019, Keer, Storch et al. 2022). Furthermore, the significant increase in *lepa* expression in hypothyroid zebrafish suggests that endocrine signaling axes may synergize, further expanding the array of potential adult morphologies attainable along a shared axis of change.

In anadromous salmonids, smoltification in preparing for migration out to sea requires orchestration of hormonal and physiological switches. A hypothesis stemming from lacustrine populations is that selective pressure on the ancestral riverine charr population was relaxed upon colonization of the lake, as the newly resident population of charr became obligate freshwater residents. Such an initial shift in developmental programs may constitute a common node among lacustrine-locked charr, biasing the direction of adaptation to generate similar forms among independent lineages, and thereby laying the foundation upon which more trophically specialized morphologies may arise. In this context, exclusion of the smoltification stage from the charr life cycle may permit lacustrine adaptive diversification. Endocrine signaling pathways, including the growth, hypothalamic-pituitary-interrenal, and thyroid axes drive physiological, morphological and behavioral changes during smoltification. In landlocked salmonid populations, the growth and hypothalamic-pituitary-interrenal axes, which ordinarily induce physiological changes for a transition to seawater, are upregulated, while the thyroid axis likely maintains its developmental, physiological, and adaptive significance (McCormick, Regish et al. 2019). In support of this model, thyroid hormone signaling is selectively modified during freshwater colonization and subsequent adaptive radiations of the threespine stickleback, *Gasterosteus aculeatus* (Kitano, Lema et al. 2010).

The lineage-specific pattern of highly differentiating loci identified in Nosed morphs, suggests that an initial developmental state of extensive modifications to thyroid signaling and craniofacial development shared among all lineages, was further refined in these lineages. The fixation of variation in *leptina2* and a predicted *otx2b* regulatory region found in Nosed morphs over other lake groups suggests further modulation of the thyroid signaling in these lineages. The data are supported by findings of altered T_3_ levels within these lineages as adults and the presence of *mc4r* within an interval of excess allele sharing with the Bigmouth morph. Thus, beginning with an initial suite of shared genetic variants, lineage-specific, secondary elaborations may have accumulated and further catalyzed the exceptional species flock diversification in Lake Kronotskoe.

Such repeated and parallel derivations of morphotypes across the *Salvelinus* complex suggest that there is an underlying genetic framework biasing the radiations of resident lacustrine populations. Our data suggest that shared modulation of the thyroid signaling axis in tandem with craniofacial regulators may enforce such biases. The patterns of variation identified in the Lake Kronotskoe radiation point to a fundamental genetic groundwork for craniofacial evolution and a common axis for morphological change.

## Methods

### Field material collection

Charrs that passes the spawning season, adults without spawning changes in color and head and reproductive states, were sampled in Lake Kronotskoe. Adult riverine Dolly Varden charrs were collected in the nearest watercourse draining the opposing slope of Valaginskiy range. Blood, pectoral fin tissue and pictures were collected. Blood for thyroid hormone test was carefully collected from the caudal vessel with a Vacuette serum tube. The distal part of the right pectoral fin (finclip, 0.2-0.3 cm^2^) was taken with scissors and fixed in pure alcohol for DNA analysis. Fish were photographed, treated with the antibacterial and antifungal solution (Melafix and Pimafix, API) for 30 min, and released if the fish did not display any signs of injury and/or infection in 48 hrs. All catches were carried out in accordance with the Russian Federal Act on Specially Protected Natural Areas (OOPT) N33-ФЗ 14/03/1995, article 10, and Plan of the research activities of Kronotsky Nature Reserve. The procedures with fish were approved by the Bioethics Commission of the AN Severtsov Institute of Ecology and Evolution, Russian Academy of Science.

### Phylochip targeted sequence enrichment design

We aimed to create a pan-Salmoniformes targeted sequence capture design that can enrich sequencing libraries for conserved genetic regions across a broad diversity of available salmon genomes. This design targets protein-coding exons as well as a set of conserved non-protein coding elements (CNEs), miRNA hairpins, and ultraconservative non-coding elements (UCNEs). The majority of capture baits were derived from the Atlantic salmon genome (*Salmo salar*, ICSASG_v2)(Davidson, Koop et al. 2010), with inclusion of regions from the genome of rainbow trout (*Oncorhynchus mykiss*, AUL_PRJEB4421_v1)(Berthelot, Brunet et al. 2014) that were either not represented in the Atlantic salmon genome or were <85% identity to a capture target within the rainbow trout genome. As these fish bracket both sides of the salmon phylogeny (**Supp. Fig. 1A**), the ‘Phylochip’ design strategy enables DNA from the majority of salmonids target regions to be efficiently enriched using this one capture design.

As the Atlantic salmon genome was not annotated at the time of capture design, annotated coding sequences were isolated from the rainbow trout and northern pike (*Esox lucius*, GCF_000721915.2_ASM72191v2)(Rondeau, Minkley et al. 2014) genomes. These were then identified within the Atlantic salmon genome via BLASTN (ncbi-blast-2.2.30+; parameters ‘-max_target_seqs 1 -outfmt 6’), and these hits used in the capture design. Genes from rainbow trout that were not identified in Atlantic salmon or that had <85% identity to the best BLAST hit within the Atlantic salmon genome were also retained in the capture design. CNEs were defined from the constrained regions in the Ensembl compara 11-way teleost alignment (Ensembl release-84)(Herrero, Muffato et al. 2016). To reserve space in the capture design, only CNEs ≥75bp in length were included in the capture baits. These CNEs were extracted from the Japanese medaka (*Oryzias latipes*, MEDAKA1), three-spined stickleback (*Gasterosteus aculeatus*, BRAOD S1), and zebrafish (*Danio rerio*, GRCz10.84) genomes using Bedtools (v2.23.0) intersectBed (Quinlan and Hall 2010). miRNA hairpins were extracted from miRbase and ultraconservative elements (UCNEs) from UCNEbase (Kozomara and Griffiths-Jones 2010, Dimitrieva and Bucher 2013). As with protein coding exons, these elements were identified within each reference genome using BLASTN (ncbi-blast-2.2.30+; parameters ‘-max_target_seqs 1 -outfmt 6’). miRNA hairpins were padded to be at least 100 bp to improve capture specificity.

From these targeted regions, the specific SeqCap EZ Developer (cat #06471684001) oligonucleotide capture baits were made in collaboration with the Nimblegen design team. Capture baits are strategically designed to standardize oligo annealing temperature, remove low complexity DNA regions and to reduce the oligo sequence redundancy. The capture design targeted sequence from 558,882 genomic regions (97,049,118 total bp) across the two salmonid genomes. This included including 460,210 protein coding exons, 93,973 CNEs, 1,082 miRNAs and 3,617 UCNEs (**Supp. Fig. 1**).

### DNA extraction and preparation of sequencing libraries

Tissue from finclips was digested and genomic DNA was column purified using QIAGEN DNeasy Blood & Tissue Kit (QIAGEN 69506). Genomic DNA was extracted from finclips of 1 *S. leucomae*nis, 3 riverine Dolly Varden charr, 8 Bigmouth morphs, 10 Longhead morphs, 5 Nosed1 morphs, 7 Nosed2 morphs, 5 Nosed3 morphs, 6 Smallmouth morphs, and 6 White morphs. Pools of genomic DNA were produced for each lineage such that genomic DNA from every individual in a lineage pool was equally represented. The pooled samples were sheared to a target size of 200bp in Tris-HCl EDTA shearing buffer (10mM Tris-HCl, 0.1mM EDTA, pH 8.0). Mechanical shearing was performed using a Covaris E220 ultrasonicator (duty cycle, 10%; intensity, 5; cycles/burst, 200; time, 400 seconds; temperature, 8°C) and Covaris microTUBE Snap-Cap tubes (Covaris 520045). Sequencing libraries were produced using the KAPA HyperPrep Kit (Roche 07137923001) using 500ng of starting material for each library. Library preparation was conducted by following the SeqCap EZ HyperCap Workflow Version 1.2. The sequencing library for the Nosed2 samples utilized enzymatic shearing using the KAPA HyperPlus Kit (Roche 07962401001) and 100ng of starting material (SeqCap EZ HyperCap Workflow Version 3.0). Fragment size and DNA concentration were quantified using Agilent 2100 BioAnalyzer and High Sensitivity DNA Chips (Agilent 5067-4626). Paired-end, 150bp Illumina HiSeq sequencing was performed on a pool consisting of multiple barcoded libraries.

### Trimming Adapters and Read Mapping

Illumina adapter sequences were removed from reads using Trimmomatic v0.36 (Bolger, Lohse et al. 2014). Trimmed and masked reads were aligned to the *Salvelinus alpinus* reference genome (RefSeq Assembly accession: GCF_002910315.2) using NextGenMap v0.5.5 (Sedlazeck, Rescheneder et al. 2013). The flag - - strata 1 was so only the highest scoring read mappings were recorded in the alignment file.

### Variant Calling and Filtering

The variants used to reconstruct the phylogeny and to conduct the principal components analyses were derived from sample of *S. leucomaenis*, anadromous Dolly Varden, and all 7 members of the species flock. samtools v1.15.1 was used to fix mates, mark duplicates, and filter reads below the minimum mapping quality set to -q 30 (Danecek, Bonfield et al. 2021). bcftools v1.13 was used to call and filter variants (Danecek, Bonfield et al. 2021). Only the variants with quality scores >= 20, depth of coverage on a per-sample basis between 10 and 500 reads, fraction of missing genotypes F_MISSING <= 0.72. SNPs within 2bp of indels and other variant types were excluded, and minor allele frequency > 0.05. The quality filtered VCF file contained 623,619 variants. The set of variants used for introgression analyses, quantification of Fst, π, and GOterm enrichment analyses were called and filtered from alignments of the anadromous Dolly Varden, Bigmouth, Longhead, Nosed 1, Nosed 3, Smallmouth, and White lineages (*S. leucomaenis*, and Nosed 2 were excluded). The calling and filtering criteria are identical to the conditions described above except for the depth thresholds. Those were filtered on a per site basis for coverage between 70 and 3500 reads. This VCF file contained 526,811 variants.

### Coverage of Targeted elements

Coverage statistics were derived using the BEDTools v2.21.1 coverage function (Quinlan and Hall 2010). Alignment files were intersected with a bed file containing the positions of each targeted element. From this intersection, the average depth of coverage was quantified per base.

### Deriving Phylogeny

The phylogeny was derived using IQ-TREE v1.6.12 (Nguyen, Schmidt et al. 2015). The input consisted of 622,831 nucleotide sites, including 22,701 parsimony informative variants. The ModelFinder function (Kalyaanamoorthy, Minh et al. 2017) determined the base substitution model of best fit to be a transversion model with empirical base frequencies and a proportion of invariable sites (TVM + F + I). 1,000 ultrafast bootstrap replicates (Hoang, Chernomor et al. 2017) quantified support for the phylogeny.

### Principal Component Analysis of Sequence Variation

PLINK v1.90b7 was utilized to conduct principal component analysis (Purcell, Neale et al. 2007). Linkage pruning was conducted using 50kb windows, 10bp step size, and R^2^ > 0.1. The linkage pruned dataset consisted of 65,488 variant sites. The PLINK eigenvector and eigenvalue outputs were plotted in R.

### Introgression Analysis

Introgression was quantified for all trios in the phylogeny using Dsuite v0.4 (Malinsky, Matschiner et al. 2021) Dtrios. Riverine Dolly Varden was specified as the outgroup. The f-branch statistic was depicted as a matrix by taking the output from Dsuite Fbranch and running the Dsuite dtools.py script to generate a plot. Dinvestigate was used to generate sliding windows of 40 variants per window and 50% overlap.

### Calculating pairwise Fst and Tajima’s Pi

To quantify genetic differentiation, pairwise Fst was calculated using PoPoolation2 v1201 (Kofler, Pandey et al. 2011). The software package allows Fst to be quantified in sliding windows or in a genewise manner. Sequencing alignment data were converted into the mpileup format using SAMtools v1.13 (Li, Handsaker et al. 2009). The PoPoolation2 program mpileup2sync (-- min-qual 20) generated the sync file used as input to calculate Fst in sliding windows. Popoolation2 calculated Fst based on allele frequency (Hartl, Clark et al. 1997). The PoPoolation2 program fst-sliding (--min-coverage 20 --min-count 3 --max-coverage 200) was used to calculate Fst in non-overlapping sliding windows. To assess which genes were most differentiating between populations, the function create-genewise-sync was utilized to intersect the sync file with a gtf containing all targeted regions in the *S. alpinus* genome, and filtered according to the same depth and minor allele count criteria as sliding windows analyses. Prior to filtering for depth, Fst was calculated for 59,478 genes and 22,590 CNEs. Noncoding elements were associated with putative regulatory targets by following the GREAT workflow to establish basal regulatory windows. A BED file of CNE loci was intersected with Intervals spanning 5kb upstream of and 1kb downstream from transcriptional start sites, with up to a 1 Mb extension (McLean, Bristor et al. 2010). To quantify nucleotide diversity, Tajima’s Pi was calculated using PoPoolation v1.2.2 (Kofler, Orozco-terWengel et al. 2011). The software also enables quantification of Tajima’s Pi in sliding windows. The same depth criteria that were used for Fst sliding window quantifications were used to calculate Tajima’s Pi in sliding windows for individual lineages.

### ELISA for thyroid hormone in blood

Serum samples were transferred to 2 ml specimen tubes and centrifuged at 12 000 g for 10 min with Eppendorf MiniSpin. Serum was then collected into Eppendorf 1.5 ml tubes and placed in in freezer at 24-26C. The total triiodothyronine (T3, bioactive form of thyroid hormone) concentration in plasma was evaluated by enzyme-linked immunosorbent assay (Monobind Total Triiodothyronine (tT3) test system, Monobind Inc, USA) and measured the hormone in accordance with the manufacture protocol using StatFax 303 Plus strip reader (Awareness Technology Inc, USA).

### Zebrafish thyroid follicle ablations

*Danio rerio* were all of the line *Tg(tg:nVenus-v2a-nfnB)* (McMenamin, Bain et al. 2014). Briefly, clutches of transgenic embryos were sorted for nVenus expression at 4 dpf then treated overnight with either 1% DMSO (for control euthyroid fish) or with 1% DMSO and 10 mM metronidazole, which induces conditional thyroid ablation in the *nfnB*-expressing thyroid follicle cells. Thyroid ablation was visually confirmed at 5dpf.

### Quantification of maxilla position

AMIRA (version 6.0.0) was used to visualize μCT scans of adult zebrafish skulls (Blythe, Nguyen et al. 2022, Nguyen, Lanni et al. 2022). A line was drawn intersecting the parasphenoid at its proximal- and distal-most points to approximate the long-axis of the body. A perpendicular line was drawn from the dorsal-most position of the maxilla to intersect the parasphenoid axis. For each individual, this distance was normalized to the standard length. Samples were selected to be roughly equivalent in size with standard lengths ranging from 19.0mm to 20.5mm. n = 10 DMSO control individuals and n = 11 MTZ treated individuals.

### Gene expression quantification

Euthyroid controls and hypothyroid siblings were decapitated at 7 or 14 dpf posterior to the operculae, and three sets of 20 heads for each condition were stored in RNA*later*™ Stabilization Solution (Thermo Fisher, Waltham MA, USA) at −20°C. RNA was extracted using a *Quick-*RNA™ Microprep Kit (Thermo Fisher, Waltham MA, USA) and cDNA libraries synthetized using SuperScript™ IV Reverse Transcriptase (Thermo Fisher, Waltham MA, USA). Using primer sequences (**Table 1**) for *actinB1*, *lepa*, *otx2b*, *pax1b*, *pax1a*, *sf3b4*, and *slc26a10*, qPCR was performed with PowerUp™ SYBR™ Green Master Mix (Thermo Fisher, Waltham MA, USA) on a QuantStudio™ 3 Real-Time PCR System (Thermo Fisher, Waltham MA, USA) with three technical and biological replicates. Results were analyzed using DataConnect Software. Relative gene expression was calculated using the ΛλΛλCT method with *actinB1* serving as the housekeeping gene (Livak and Schmittgen 2001).

**Table 1.**
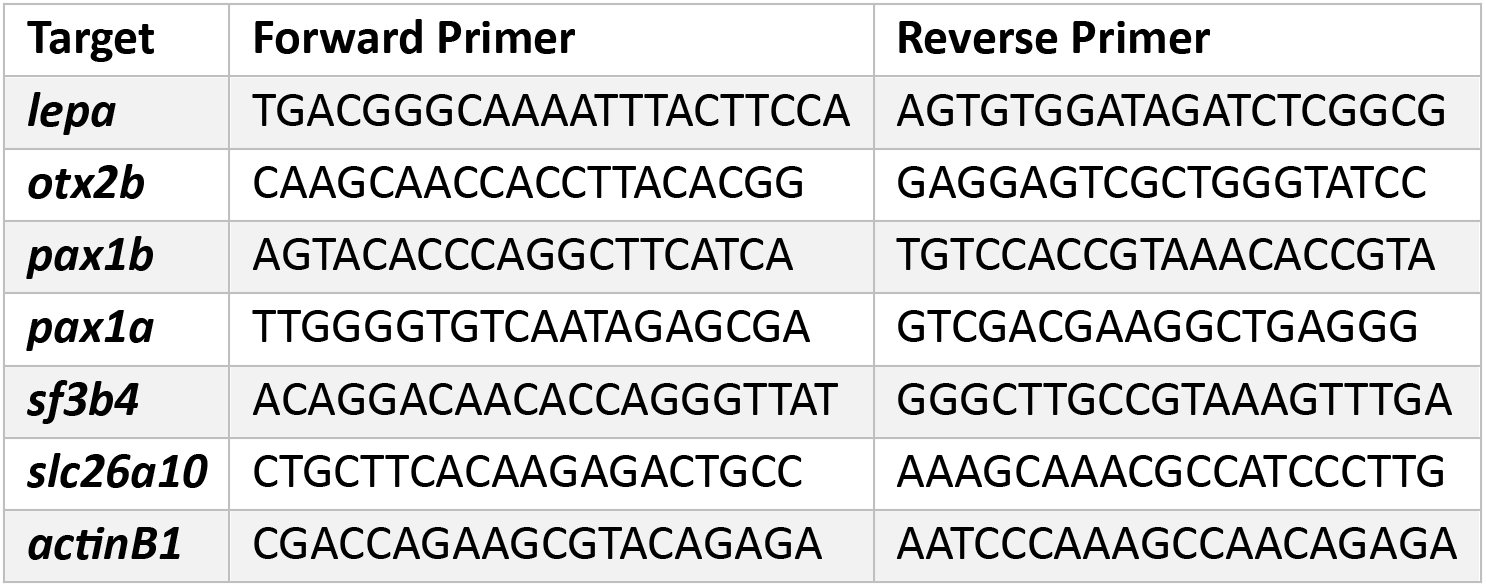
Sequences for primers used in qPCR.

### Geometric morphometrics

Pictures of dry osteological samples (14-27 of each morph) were used for landmarking (Saltykova, Markevich et al. 2015). Using TPSdig v2.0 (Rohlf 2015), we digitized landmarks (LM), most of which have been used for the homologous bones of zebrafish (Keer, Storch et al. 2022) and charrs (Guðbjörg Ósk, Laura-Marie von et al. 2024): six LM for dentary, ten LM for anguloarticulare; eleven LM for hyomandibula; and eight LM for parasphenoid (**Figure 7 – figure supplement 1**). Shape analysis was performed in MorphoJ v1.06d (Klingenberg, 2008). We implemented Generalized Procrustes superimposition and assessed variation in the shape with Principal component analysis (PCA). For better visualization of shape variability along PC1/PC2, we created a wireframe mesh connecting landmarks. To estimate the shape differences between the morphs, we implemented the Procrustes ANOVA and Canonical Variate (CV) analysis with a calculation of pairwise Procrustes distances (10,000 permutation rounds).

## Acknowledgements

We owe thanks to Dr. Stacy Nguyen for providing uCT data and to Dr. E.A. Saltykova for providing images of dry osteological samples.

## Data availability

We will make relevant sequencing files available on Zenodo.

---

**Figure 2 – Supplementary Figure 1.**
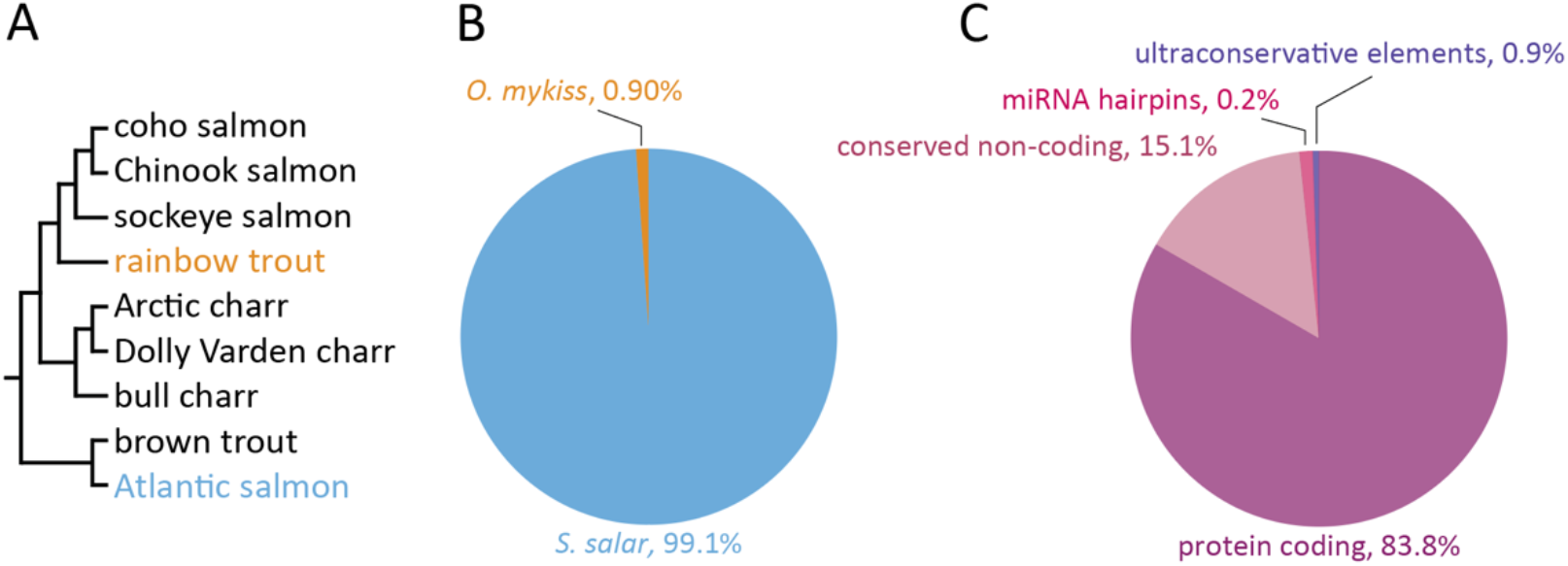
Design of pan-Salmoniformes targeted capture array. **(A)** Conserved bait sequences were derived from Atlantic salmon (*Salmo salar*) and rainbow trout (*Oncorhynchus mykiss*) reference genomes. **(B)** Relative contributions of *S. salar* and *O. mykiss* derived bait sequences to the capture array. **(C)** Classifications and relative abundances of conserved elements targeted for capture.

**Figure 2 – Supplementary Figure 2.**
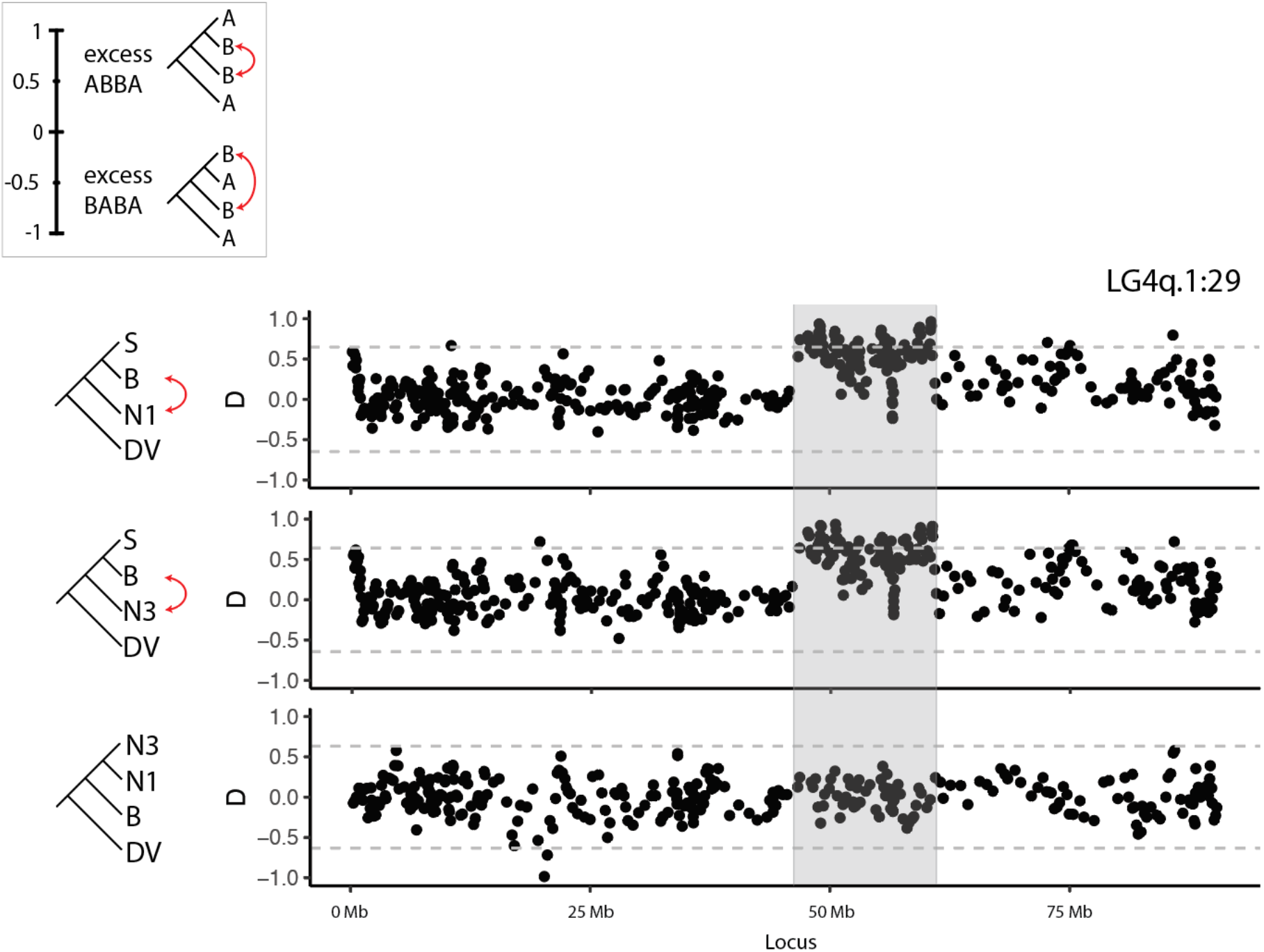
Sliding window plots of D showing a *large* interval of excess allele sharing. Sliding window plots for trios consisting of Smallmouth (S), Bigmouth (B), and Nosed 1 (N1), of S, B, and Nosed 3 (N3), and of N3, N1, and B. Positive D values indicate an excess of the ABBA pattern (red arrows), while negative values indicate an excess of the BABA pattern. The three plots show a common pattern of excess of allele sharing overlapping with between B and N1 and B and N3, while there is no excess of allele sharing between B and N1 over B and N3. Horizontal lines signify 3SDs from the mean.

**Figure 2 – Supplementary Table 1.** Summary of reads aligned and targeted element coverage.

| Morphotype | No. Individuals<br>Pooled | No. Reads<br>(million) | No. Reads<br>Mapped | Median<br>Depth | Mean<br>Depth | "Conservome"<br>Coverage |  |
| --- | --- | --- | --- | --- | --- | --- | --- |
|  |  |  |  |  |  | 2x | 10x |
| Bigmouth | 8 | 72.3 | 35.3 | 66 reads | 88 reads | 91.6% | 88.5% |
| Dolly Varden | 3 | 64.1 | 32.4 | 61 reads | 82 reads | 91.6% | 88.3% |
| Longhead | 10 | 46.1 | 22.8 | 41 reads | 56 reads | 91.1% | 85.7% |
| Nosed 1 | 5 | 68.8 | 34.9 | 65 reads | 87 reads | 91.7% | 88.5% |
| Nosed 2 | 7 | 49.5 | 17.5 | 16 reads | 20 reads | 88.7% | 65.5% |
| Nosed 3 | 5 | 68.4 | 34.1 | 63 reads | 84 reads | 91.6% | 88.5% |
| Smallmouth | 6 | 43.1 | 21.1 | 38 reads | 52 reads | 90.8% | 84.9% |
| *S. leucomaenis* | 1 | 55.9 | 25.5 | 50 reads | 67 reads | 90.5% | 85.9% |
| White | 6 | 36.9 | 18.5 | 34 reads | 46 reads | 90.6% | 83.8% |

**Figure 2 – Supplementary Table 2.** Table of significantly underrepresented GO terms. This table contains the set of significantly underrepresented GO terms among all sample libraries (Benjamini-Hochberg FDR < 0.05).

| GO name | GO accession | bin size |
| --- | --- | --- |
| activation of MAPK activity | GO:0000187 | 15 |
| anaphase-promoting complex | GO:0005680 | 51 |
| anaphase-promoting complex-dependent catabolic process | GO:0031145 | 41 |
| ATP-dependent chromatin remodeling | GO:0043044 | 133 |
| ATPase activity | GO:0016887 | 262 |
| calcium ion binding | GO:0005509 | 1185 |
| calcium, potassium:sodium antiporter activity | GO:0008273 | 52 |
| carbohydrate binding | GO:0030246 | 111 |
| carbohydrate metabolic process | GO:0005975 | 204 |
| cell adhesion | GO:0007155 | 541 |
| chemokine activity | GO:0008009 | 72 |
| cullin-RING ubiquitin ligase complex | GO:0031461 | 23 |
| DNA binding | GO:0003677 | 2972 |
| DNA integration | GO:0015074 | 1571 |
| DNA-templated transcription, initiation | GO:0006352 | 36 |
| ferric iron binding | GO:0008199 | 24 |
| fructose-bisphosphate aldolase activity | GO:0004332 | 27 |
| hexose metabolic process | GO:0019318 | 34 |
| homophilic cell adhesion via plasma membrane adhesion molecules | GO:0007156 | 362 |
| iron ion transport | GO:0006826 | 23 |
| isomerase activity | GO:0016853 | 45 |
| MAP kinase activity | GO:0004707 | 52 |
| nucleic acid binding | GO:0003676 | 2731 |
| oligopeptide transport | GO:0006857 | 20 |
| phosphatidylinositol metabolic process | GO:0046488 | 49 |
| phosphatidylinositol phosphate kinase activity | GO:0016307 | 47 |
| receptor tyrosine kinase binding | GO:0030971 | 28 |
| regulation of mitotic metaphase/anaphase transition | GO:0030071 | 44 |
| sensory perception of sound | GO:0007605 | 69 |
| skeletal muscle fiber development | GO:0048741 | 44 |
| SWI/SNF complex | GO:0016514 | 147 |
| transcription coactivator activity | GO:0003713 | 164 |
| transcription factor TFIID complex | GO:0005669 | 38 |
| transmembrane transport | GO:0055085 | 1148 |
| transmembrane transporter activity | GO:0022857 | 306 |
| transporter activity | GO:0005215 | 218 |
| transposition, DNA-mediated | GO:0006313 | 1551 |
| ubiquitin protein ligase binding | GO:0031625 | 28 |
| vesicle-mediated transport | GO:0016192 | 250 |

**Figure 2 – Supplementary Table 3.** Table of D_tree_ scores, ƒ _4_-admixture ratios, and Z-scores for each of the 20 trios contained within the Lake Kronotskoe species flock. 16 trios were found to have a significant, though minimal, contribution of introgressed alleles (asterisks)(Holm-Bonferoni, FWER < 0.01). B, Bigmouth; L, Longhead; N1, Nosed 1; N3, Nosed 3; S, Smallmouth; W, White.

| Pop 1 | Pop 2 | Pop 3 | D<sub>tree</sub> (%) | f<sub>4</sub>-ratio (%) | p-value | Z-score |
| --- | --- | --- | --- | --- | --- | --- |
| S | N3 | W | 3.9 | 8.6 | 0 | 13.0* |
| S | N1 | W | 3.7 | 8.2 | 0 | 11.4* |
| S | N3 | L | 2.8 | 6.3 | 0 | 11.4* |
| S | N1 | L | 2.2 | 4.9 | 1.18E-16 | 8.3* |
| S | B | N1 | 4.2 | 9.0 | 1.56E-15 | 8.0* |
| S | B | W | 2.2 | 4.7 | 1.26E-14 | 7.7* |
| B | N3 | W | 1.8 | 4.1 | 2.44E-14 | 7.6* |
| L | W | N1 | 2.2 | 4.5 | 2.73E-13 | 7.3* |
| L | W | B | 1.7 | 3.9 | 2.76E-13 | 7.3* |
| B | N1 | W | 1.7 | 3.7 | 2.11E-12 | 7.0* |
| S | B | N3 | 3.6 | 8.3 | 2.31E-11 | 6.7* |
| L | W | N3 | 1.8 | 3.8 | 4.65E-09 | 5.9* |
| B | N3 | L | 1.8 | 4.0 | 6.52E-07 | 5.0* |
| S | B | L | 1.1 | 2.4 | 1.01E-04 | 3.9* |
| B | N1 | L | 1.2 | 2.6 | 3.59E-04 | 3.6* |
| N1 | N3 | L | 0.7 | 1.5 | 1.13E-03 | 3.3* |
| L | W | S | 0.5 | 1.4 | 2.86E-03 | 3.0 |
| N1 | N3 | S | 0.4 | 0.9 | 5.72E-02 | 1.9 |
| N1 | N3 | W | 0.2 | 0.5 | 2.79E-01 | 1.1 |
| N3 | N1 | B | 0.3 | 0.7 | 2.86E-01 | 1.1 |

**Figure 7 – Supplementary Figure 1.**
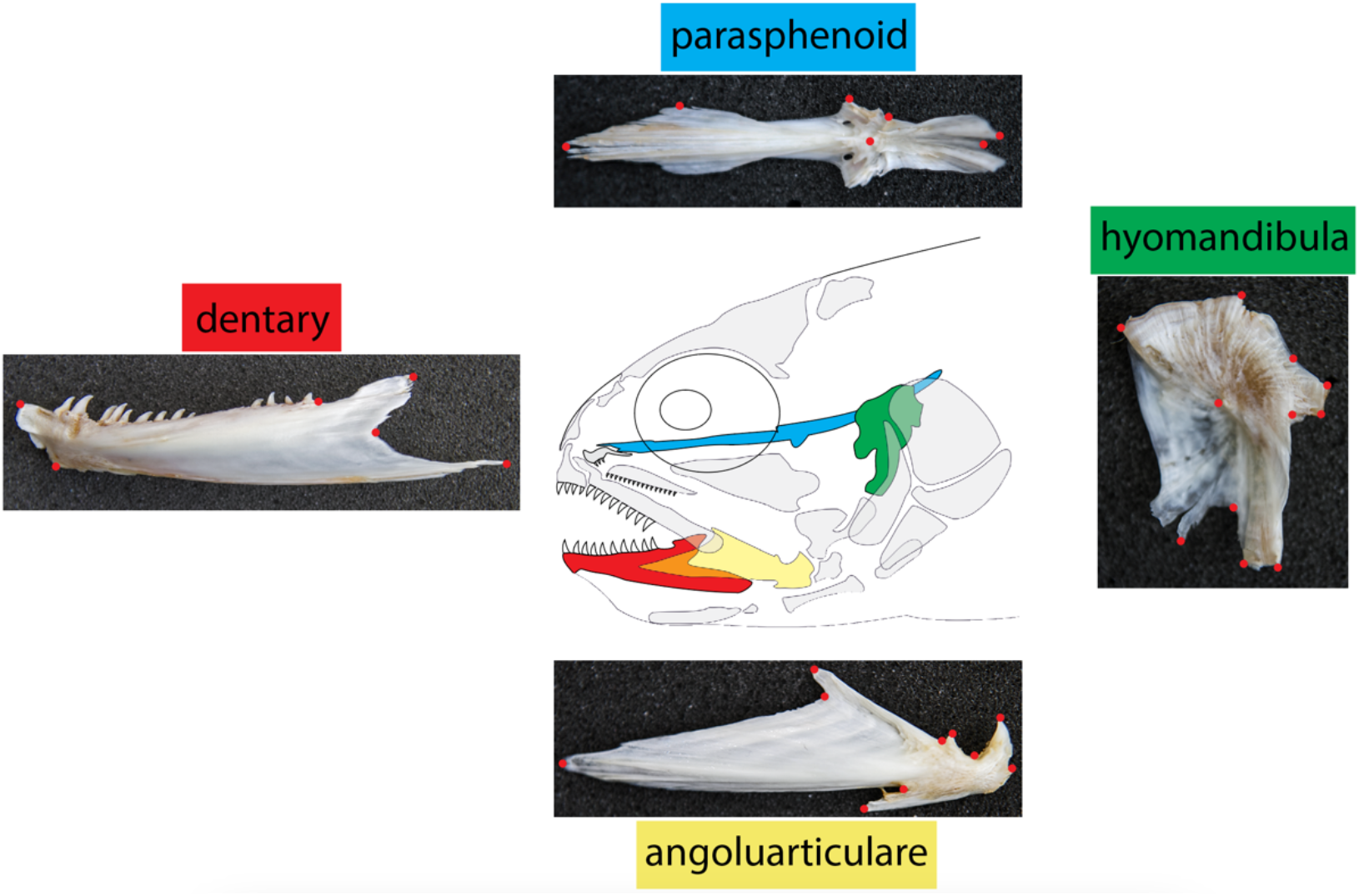
Landmarks used to geomorphic morphometric analyses. Landmarks were assigned to each of four bones. The dentary (red), the parasphenoid (blue), the hyomandibula (green), and the anguloarticulare (yellow).

**Figure 7 – Supplementary Figure 2.**
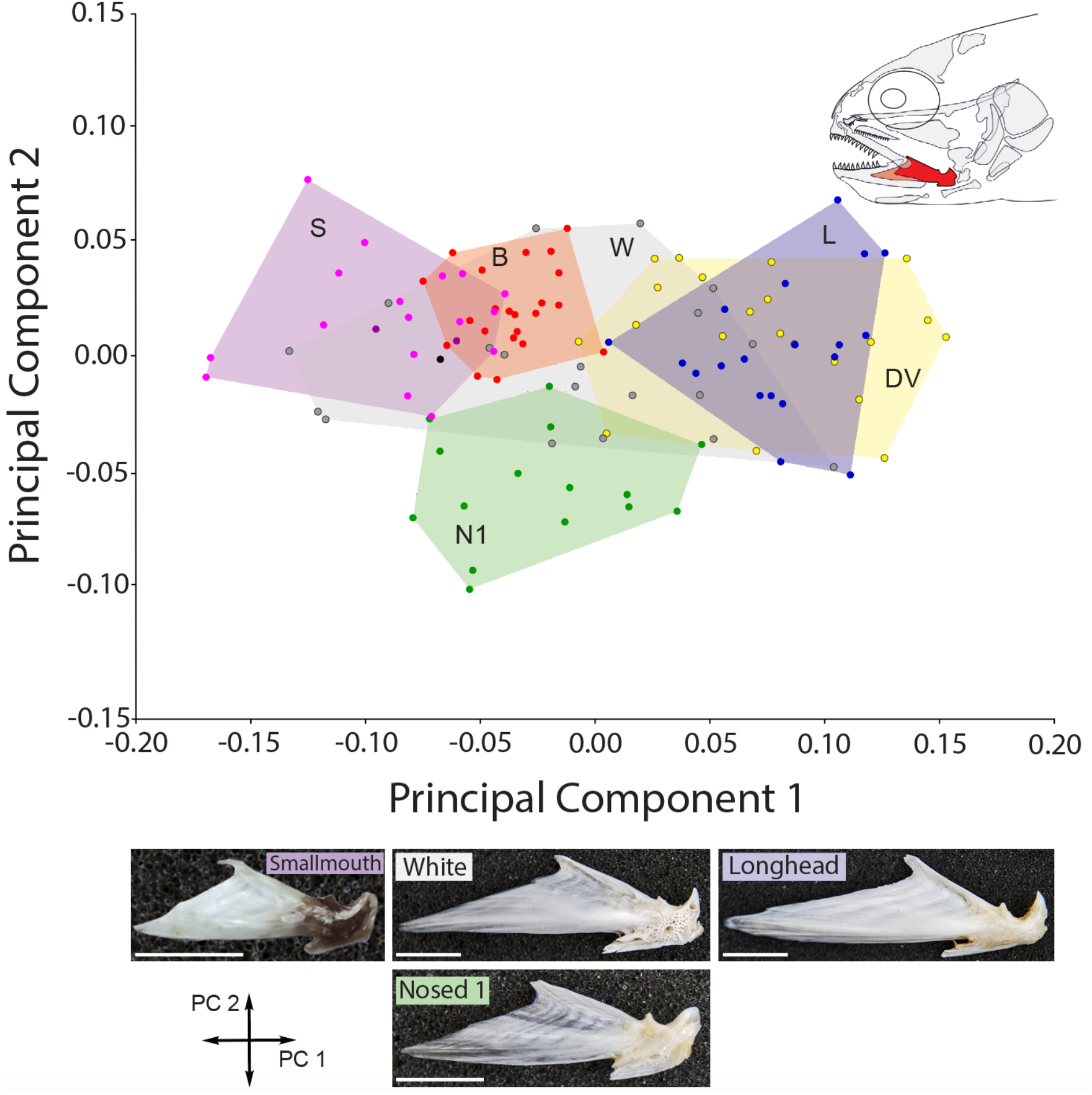
Geometric morphometric analyses of anguloarticulare among Lake Kronotskoe morphs. Procrustes analyses identified highly significant differences in shape (Procrustes ANOVA F_80;1760_=18.88 p˂0.0001). Scale bar = 10mm.

**Figure 7 – Supplementary Figure 3.**
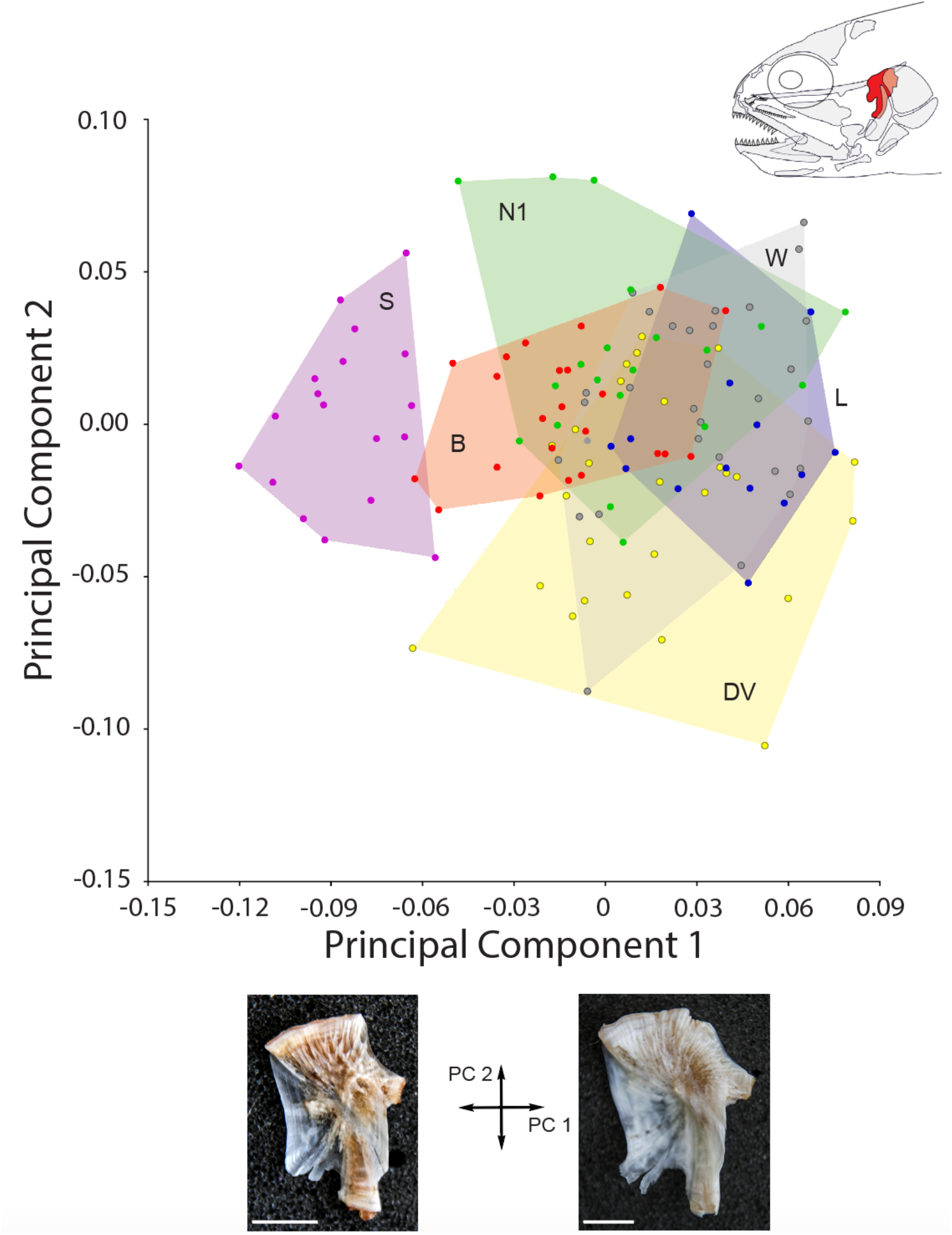
Geometric morphometric analyses of hyomandibula among Lake Kronotskoe morphs. Procrustes analyses identified significant differences in shape (Procrustes ANOVA F_60;1212_=6.55 p˂0.001) with Smallmouth morphs possessing the most distinct shape.

**Figure 7 – Supplementary Figure 4.**
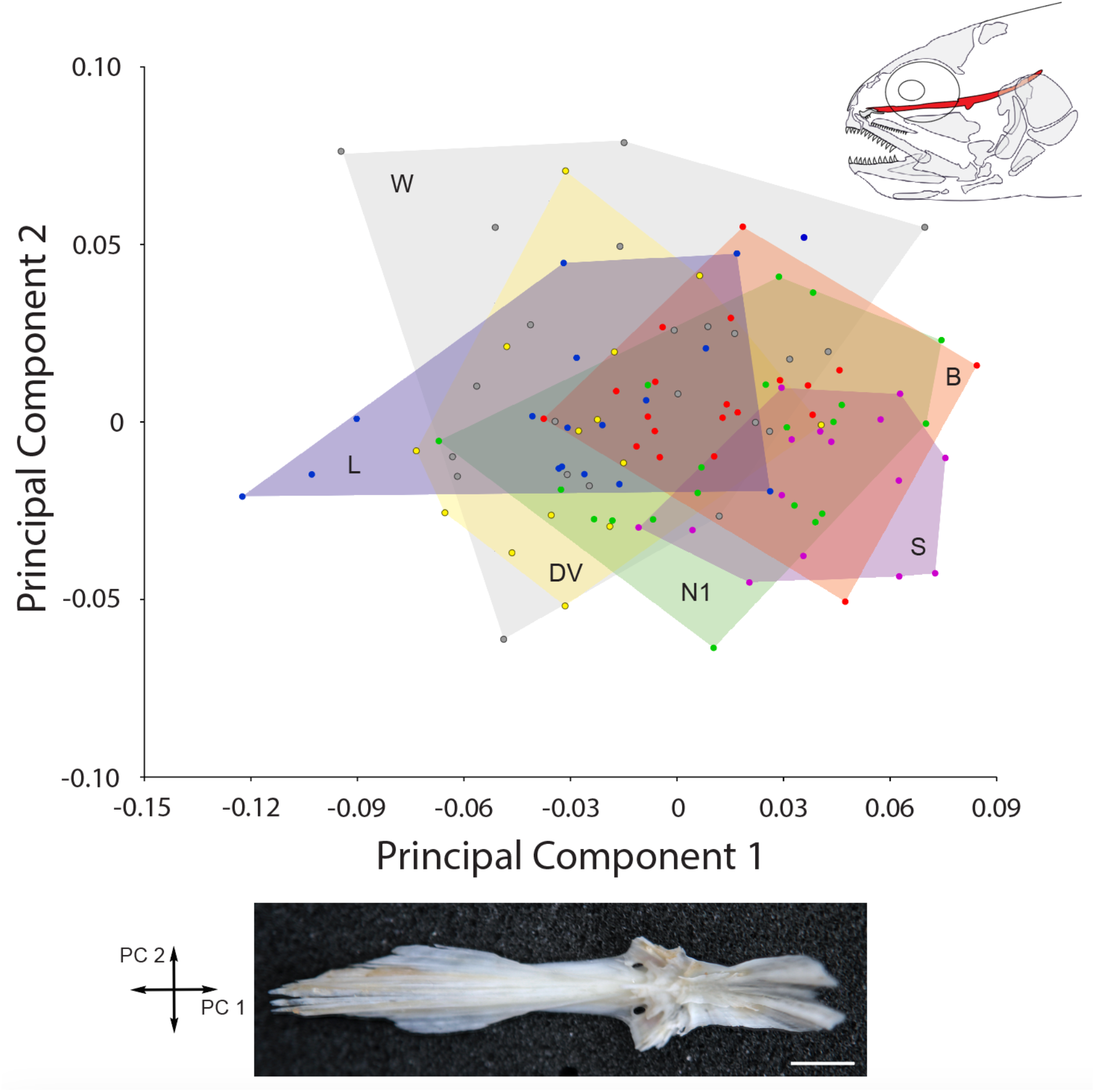
Geometric morphometric analyses of parasphenoid among Lake Kronotskoe morphs. Procrustes analyses identified no significant differences in shape. Scale bar = 10mm.

